# Hidden but discoverable diversity in the global microbiome

**DOI:** 10.1101/2025.06.26.661807

**Authors:** V Prasoodanan PK, OM Maistrenko, A Fullam, DR Mende, E Kartal, LP Coelho, A Spang, P Bork, TSB Schmidt

## Abstract

Cataloguing Earth’s biodiversity remains one of the most formidable challenges in biology, and the greatest diversity is expected to reside among the smallest organisms: microbes. Yet the ongoing census of microbial life is hampered by disparate sampling of Earth’s habitats, challenges in isolating uncultivated organisms, limited resolution in taxonomic marker gene amplicons, and incomplete recovery of metagenome-assembled genomes (MAGs). Here, we quantified discoverable bacterial and archaeal diversity in a comprehensive, curated cross-habitat dataset of 92,187 metagenomes. Clustering 502M sequences of 130 marker genes, we detected 705k bacterial and 27k archaeal species-level clades, the vast majority of which was hidden among ‘unbinned’ contigs. At deeper taxonomic levels, we estimate that 10 archaeal and 145 bacterial novel phyla and around 80k novel genera are discoverable in current data. We identified soils and aquatic environments as novel lineage recovery hotspots, yet predict that discovery will remain in full swing across habitats as more data accrues. Finally, we show that prokaryotic diversity follows power laws, confirming century-old hypotheses on clade size patterns and suggesting that novel lineages arise within common (and fractal) evolutionary patterns, comparable to those among eukaryotic clades and viruses, along the full depth and breadth of the Tree of Life.

## Introduction

Microbial life on Earth has deep evolutionary roots, and is ubiquitous and abundant in extant ecosystems: prokaryotes (that is, Bacteria and Archaea) likely emerged more than 4 billion years ago ^1^, have been detected in virtually every environment on the planet, and are estimated to account for ∼10^30^ cells and 10^16^-10^18^ g of biomass ^2,3^. This long history of evolution among large numbers of individuals across a comprehensive ecological range is reflected in a great accumulated phylogenetic diversity, vastly exceeding that of multicellular life ^4^. Yet it remains unclear just how diverse extant Bacteria and Archaea really are: estimates based on extrapolations from abundance distributions of individual samples vary by orders of magnitude, predicting anywhere between 10^6^ and 10^12^ prokaryotic species worldwide ^5–8^.

Only a fraction of this predicted diversity has been accounted for in existing sequence data. Using 16S rRNA amplicon sequences, previous studies estimated the number of species-level Operational Taxonomic Units (OTUs) in publicly available datasets at 35.5k in 2004 ^9^, 210k in 2014 ^8^, 109k in 2016 ^10^ and 740k in 2019 ^7^. Based on *rarefaction curves*, tracking the number of newly discovered types (i.e., species) as more data (i.e., samples) are added to the survey, these analyses concluded that at least in some habitats, the rate of species discovery was slowing down (reflected in a ‘flattening off’ in the rarefaction curves).

While 16S rRNA data continues to be more abundant in public repositories (e.g., the Microbe Atlas Project encompasses 1.7M amplicon samples ^11^), integrated surveys of prokaryotic diversity increasingly rely on isolate and metagenome-assembled genomes (MAGs) to overcome some of the limitations (see Supplementary Discussion) of amplicon-based datasets. In metagenomic data, clade-level diversity (e.g., species) is often defined based on sequence similarity in either a subset of near-universal taxonomic marker genes ^12–14^ or of entire genomes ^15^. For example, proGenomes3 ^16^ encompasses 41k species-level clusters of high-quality isolate genomes from NCBI’s RefSeq and GenBank (v3, 01-2023), whereas the Genome Taxonomy Database (GTDB) r226 (04-2025) delineates 143k species, combining isolate genomes and high-quality MAGs into a systematic reference taxonomy ^17^. In addition, large-scale MAG catalogues generated from publicly available data continue to add to the census, reporting e.g. 4.6k species-level genomes in the human gut microbiome ^18^, 8.3k species in the ocean ^19^, 21k species in soils ^20^, between 0.1-13k species for the various habitats in MGnify’s Genome collection ^21^, or 18k species integrated across the habitats represented in the Joint Genome Institute’s IMG/M database ^22^. We recently developed SPIRE ^23^ which provides an integrated survey of microbial diversity across 99k manually annotated metagenomes from all sampled habitats on Earth, describing 107k species-level clusters of which 92k are entirely based on MAGs (i.e., lacking a cultivated representative).

Such genome-centric surveys have greatly expanded the scope of study for microbial diversity, ecology and evolution, in particular for uncultivated taxa: in the GTDB r226, arguably the broadest available reference for prokaryotic taxonomy, just 64 out of 189 phyla and 34.5k out of 143.6k species contain cultivated representatives. In other words, 66% of currently recognized phyla and 76% of species have been described solely based on MAGs. Nevertheless, MAG recovery remains limited in terms of both precision (due to residual binning artefacts ^24^) and recall (most bins remain incomplete with current automated binning tools ^25^). Indeed, with common workflows, most metagenomically assembled contigs remain *unbinned*: in SPIRE, just 2.36Tbp (or 10%) out of a total assembly length of 23.3Tbp end up in medium or high-quality MAGs. Arguably, these unbinned contigs circumscribe a space of ‘discoverable’ microbial diversity, i.e. the reservoir of recoverable genomes that remains untapped by current workflows, potentially comprising representatives of the so-called ‘rare biosphere’ ^26^. For example, recent studies estimated this genomically unrepresented microbial diversity at around 135k species in 18k metagenomes ^27^ and 83k species in 249k metagenomes ^28^. Indeed, novel tools emerge to make these genomes accessible: the recently developed Bin Chicken workflow of targeted co-assembly and co-binning recovered genomes for 24k previously unrepresented species from just 800 sample groups ^29^.

Here, we conducted a survey of discoverable bacterial and archaeal diversity among a set of 92,187 metagenomes (representing a filtered subset of SPIRE v1; see Methods), sampled across Earth’s microbial habitats. We track the diversity represented in 120 bacterial ^13^ and 53 archaeal ^14^ marker genes originating from reference isolate genomes ^16^, MAGs ^17,23^ and the large fraction of unbinned metagenomic contigs to quantify the number of novel lineages to be incrementally discovered in each set. Using habitat-stratified rarefactions, we tally the amount of novel genomic diversity that remains to be discovered in available public sequence data and estimate the degree of novelty expected to emerge in different environments as new metagenomes are added to the survey. We track discoverable lineages that lack genomic representation across marker gene trees to estimate their phylogenetic depth and quantify how many ‘deeper’ novel clades (i.e., unrecognized phyla, classes, orders, families and genera) remain undiscovered across different habitats. Finally, we show that the size distributions of bacterial and archaeal clades (including those inferred from unbinned genes) follow power laws, in line with century-old hypotheses by Willis ^30^ and Yule ^31^, with ramifications for biodiversity theory and the study of microbial evolution. In other words, we ask how many and which novel microbial lineages are ‘hiding in plain sight’ among unbinned contigs because they are missed by current toolkits, and how many more lineages we are poised to discover as we continue to sample Earth’s habitats.

## Results

### Just 20-50% of currently metagenomically discoverable prokaryotic species are captured by genome-centric approaches

We tracked the incremental discovery of bacterial and archaeal diversity among 92,187 metagenomic samples in SPIRE ^23^ along rarefaction curves using species-level clusters of assembled taxonomic marker genes (Figure 1A; see Methods). We estimated the total discoverable diversity in the studied dataset at ∼705k bacterial and ∼27k archaeal species (see table S1 and Fig 1B); rarefaction trajectories (Extended Figure 1) and overall species count estimates (Extended Figure 2) were remarkably consistent across the different independently queried taxonomic marker genes. Just 20k bacterial and 0.8k archaeal species comprised cultivated representatives in proGenomes3, an additional 25k and 1.7k species (for a total representation of 6.4% and 8.8%) mapped to the GTDB r220, and a further 80.5k and 4.2k species were contributed by SPIRE MAGs, for a total species-level diversity representation within genome-based datasets of 17.8% among Bacteria and 24.6% among Archaea. In other words, up to ∼75-80% of our putative species-level clusters were not captured by genomes of cultivated organisms or MAGs.

**Figure 1.**
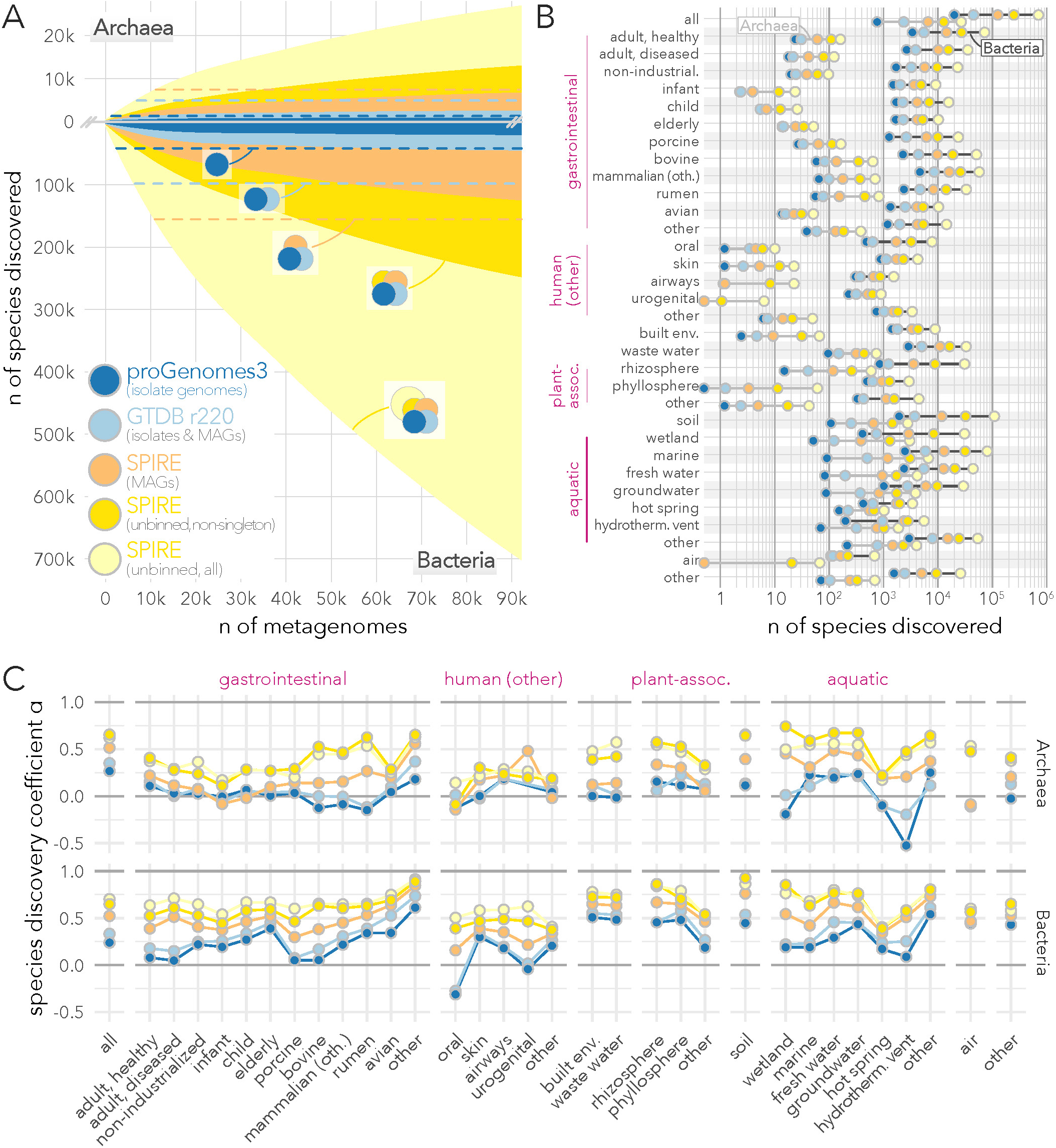
The discovery of novel prokaryotic diversity remains in full swing. **(A)** Rarefaction curves of archaeal (top) and bacterial (bottom) species discovery across ∼90k metagenomes. Based on marker gene cluster composition, we distinguished five hierarchically overlapping categories of data provenance for every inferred species (indicated as Venn diagrams): (i) those including an isolate genome represented in proGenomes3 (dark blue data series); (ii) additional species (beyond proGenomes3) represented in GTDB r220 isolate genomes and MAGs (light blue); (iii) species defined from SPIRE MAGs (orange); and species-level marker gene clusters exclusively based on unbinned contigs (i.e., not assigned to medium or high-quality MAGs), further distinguishing (iv) non-singleton clusters (containing ≥2 sequences from independent sources; dark yellow) among (v) all marker gene clusters (light yellow). Reference levels are indicated as dashed lines, i.e. the total number of species in proGenomes3, GTDB r220 and SPIRE MAGs including those *not* discoverable in metagenomes. **(B)** Total number of species-equivalent marker gene clusters per habitat; see figures S2 & S3 for corresponding habitat-stratified rarefaction curves. **(C)** Species discovery coefficients ‘α’, calculated from habitat-stratified rarefaction curves (see Methods). Values for α in [0, 1] correspond to *unsaturated* species discovery curves where additional samples continue to add novel species to the survey (analogous to ‘open’ pangenomes); lower α values indicate a more pronounced ‘flattening off’ in the rarefaction curve, indicating a more pronounced slowdown in novel species discovery; higher α values indicate a less pronounced decrease in the rate of species discovery. For α ➝ 1, species discovery is *fully unsaturated*, meaning that each newly added sample adds novel species to the survey, with no discernible ‘flattening off’ in the species discovery curve.

These ‘unbinned’ species were overrepresented among uncultivated phyla (recognized solely based on MAGs in the GTDB, with no cultivated isolates; Cohen’s d = 0.33; Wilcoxon p=0.01; Extended Figure 3A) and, compared to species with a cultivated representative, are associated with higher genomic GC content (R^2^ = 0.15, p = 1.6 * 10^−16^), but not average genome size (p = 0.338), coding density (p = 0.418) or estimated clade size (p = 0.889) in a multiple linear regression (Extended Figure 3B). Species-level cluster sizes followed a long-tailed distribution, driven by a large fraction of singleton clusters (i.e., clusters containing just one sequence; Extended Figure 4). While this is consistent with expectations on the so-called ‘rare biosphere’ of lineages with low prevalence across samples and/or low abundance within samples ^26^, a subset of singleton clusters may also arise from assembly artifacts or spurious sequences. When conservatively considering only non-singleton marker gene clusters (i.e., containing ≥2 sequences from independent sources), diversity estimates were accordingly reduced to a total of 249k bacterial and 12.8k archaeal species (Figure 1AB; table S1); this can be considered a lower bound (see Supplementary Discussion), yet still corresponded to an increase of 98% relative to the discoverable species set represented by genomes. Indeed, while around three in four unbinned marker gene clusters were singletons, they accounted for <20% of unbinned genes, with the remainder mapping to reference genomes, MAGs or larger unbinned gene clusters (Extended Figure 4). Taken together, this suggests that the vast majority of unbinned genes represent a real biological signal and likely belong to low-quality (i.e., incomplete or contaminated) MAGs, enriched in uncultivated clades.

At the same time, marker genes for around half of the species defined by (isolate) genomes in proGenomes3 and GTDB r220 were *not* represented in metagenomic assemblies (total reference dataset sizes indicated by dotted lines in figure 1A) and around 40% were singletons (Extended Figure 4). This suggests that species with cultivated representatives and those pinpointed for isolation are themselves often rare, occurring at low prevalence and/or low abundance in environmental data. Thus, the discovery gap goes both ways: isolate genomes and MAGs captured only 20-50% of discoverable species in metagenomic contigs, but metagenomic assemblies likewise failed to capture around half of the diversity represented by isolates.

### Novel species discovery remains in full swing for most of Earth’s microbial habitats

The total number of discoverable species, as well as their relative representation among cultivated organisms and MAGs, varied greatly between individual habitats (Fig 1B), with moderate correlation to sampling effort (ρ_Spearman_ = 0.51 for Bacteria and 0.09 for Archaea). In line with expectations, estimated archaeal species counts were generally 2-3 orders of magnitude below those for Bacteria in host-associated and anthropogenic habitats (including some environments with almost no detectable Archaea), roughly within 1 log in aquatic habitats and soils, and almost on par in extreme environments like hot springs and hydrothermal vents. The degree of genomic representation of discoverable species likewise varied across habitats, but was generally within one order of magnitude of the total.

We next tested whether novel species discovery showed any signs of slowing down by calculating species discovery coefficients (termed ‘α’) based on individual, habitat-stratified rarefaction curves (Fig 1C; see Methods). We observed that genome-based discovery of archaeal species was fully saturated (α ≤ 0) or approaching saturation (low positive α) in most host-associated and anthropogenic habitats. Specifically, all archaeal species represented in proGenomes3 and GTDB r220 that *can* be metagenomically discovered using current approaches, have already been accounted for in habitats like human infant gut, oral or skin samples; adding more samples to the survey is not expected to increase metagenomic representation. Similarly, anthropogenic environments (built environment and wastewater), wetlands and extreme environments (hot springs and hydrothermal vents) appear saturated in terms of metagenomic discovery of ‘known’ isolate-derived archaeal species, whereas even MAG-based archaeal species discovery showed clear signs of slowing down (low positive α). In contrast, archaeal species discovery coefficients among unbinned contigs were significantly higher across most habitats, indicating that the existing large discovery gap between binned and unbinned contigs will continue to widen, as the accumulation of unbinned novel diversity shows only moderate signs of slowing down.

Bacterial species discovery coefficients were generally higher, indicating that with current approaches, novel bacterial diversity will continue to be discovered at higher rates than for Archaea. Interestingly, the human mouth, urogenital tract and adult gut, as well as porcine and bovine gut environments are the only tested habitats in which the metagenomic discovery of isolate-represented diversity is approaching saturation. Yet even in these habitats, as well as across all others, MAG-based species discovery is expected to continue at high rates, outpaced only by species discovery among unbinned contigs (α ≥ 0.5 across most habitats). Soils, wetlands, the rhizosphere, freshwater habitats and the GI tracts of non-mammalian, non-avian animals stood out as particular hotspots of untapped diversity, with virtually no signs of a slowdown in species discovery (α ≥ 0.8). Overall, the significant difference in species discovery coefficients between reference genome sets, MAGs and unbinned contigs suggests a widening gap between *discovered* (i.e., genomically represented) and *discoverable* (in unbinned contigs) diversity, as the latter will continue to outpace the former.

### Soils and aquatic habitats remain major reservoirs of unexplored microbial diversity

Beyond species discovery rates within individual habitats, we explored how much discoverable diversity each habitat contributed to the total survey. We tracked habitat-stratified species accumulation curves – quantifying how much diversity each sequentially added habitat contributed to the survey *incrementally*, beyond what was already contained in other habitats – which revealed striking discrepancies between sampling effort and observed unique diversity.

Although our dataset contained 53,949 gastrointestinal metagenomes (corresponding to 59% of the total), these accounted for just ∼253k (or 36%) of discoverable species for Bacteria and ∼2k (or 7%) for Archaea (Extended Figure 5). This corresponds to an average of 4.7 newly added bacterial and 0.04 archaeal species per sample. Indeed, the human gut alone (across age groups, geography and disease states, yet still representing just one host species) accounted for 49% of total samples, but just 16% of discoverable bacterial and 0.8% of archaeal diversity. A further 12,098 metagenomes from non-intestinal human body sites and the built environment increased the tally to ∼72% of total samples, but contributed only an additional 13k (∼2% of the total, or 1.05 per sample) bacterial and 0.2k (<1%, or 0.02 per sample) archaeal species. In other words, while human-associated, gastrointestinal and built environment metagenomes represented almost three quarters of sampling effort, they only accounted for 38% of discoverable bacterial and 8% of archaeal diversity.

In contrast, plant-associated habitats represented just 1.5k samples (1.7% of the total), but accounted for 47k (6.7%) bacterial species discoverable beyond those contained in animal-associated and anthropogenic habitats, with the rhizosphere standing out as particular hotspot of bacterial diversity at 74.4 new species added per sample. Yet the largest steps in the collector’s curve were observed for soils (∼6% samples contributing 16% of unique bacterial and 9% of archaeal diversity, at 21.8 and 0.45 added species per sample) and aquatic habitats (∼16% of samples contributing 32% of bacterial and 78% of archaeal diversity, at 15.1 and 1.4 novel species per sample). Wetlands, including e.g. salt marshes and peatlands, were further discovery hotspots for both Bacteria and Archaea (at 53.7 and 5.0 uniquely added species per sample, respectively), as were hydrothermal vents (21.5 and 6.2 species per sample). Although these and other comparatively undersampled habitats generally contributed more novelty per sample than well-represented environments (like the human gut), the overall correlation between sampling effort and newly discovered species was only moderately negative (ρ_Spearman_ = −0.21 for Bacteria and −0.26 for Archaea).

### Uncaptured deeply branching lineages may substantially enrich the Tree of Life

We next explored how these trends in species discovery translated to deeper taxonomic levels. We inferred gene phylogenies for each of the considered 53 archaeal and 122 bacterial markers, estimated the number of represented genus-, family-, order-, class- and phylum-level clades by cutting each tree at level-specific relative evolutionary divergence (RED, ^32^) cutoffs, and summarised the resulting clade counts across a filtered subset of phylogenies (see Methods). As shown in Figure 2 for archaeal marker DNA-directed RNA polymerase subunit E (*rpoE* [TIGR00448]) as a representative example, unbinned sequences added considerable tip-level (i.e., approximately species-level) diversity for most recognized clades, while often also providing relevant ecological context via the origin of underlying samples. For example, for the family *Nitrosopumilaceae* (within the Thermoproteota phylum or TACK superphylum ^33^) that encompasses several known ammonia-oxidizing lineages, we observed a significant expansion of covered diversity among marine genera (e.g. *Nitrosopelagicus* or *Nitrosopumilus*), while linking other subclades to wetland, groundwater, soil and plant-associated habitats, supported across several phylogenies (based on different marker genes, i.e. beyond the rpoE tree in Figure 2).

**Figure 2.**
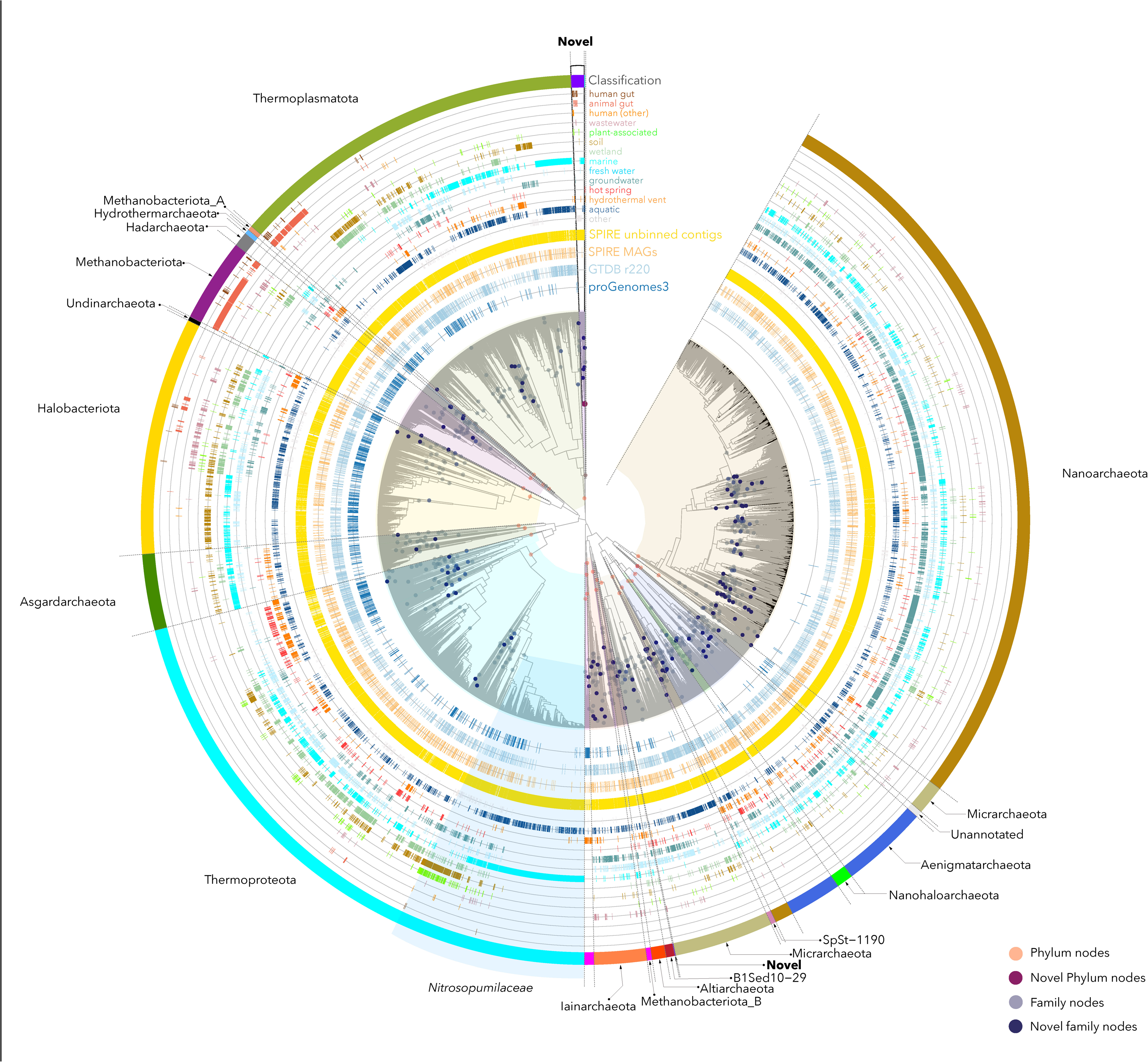
Unbinned contigs flesh out the Tree of Life among both known and deep novel clades. Example phylogeny of the archaeal RNA polymerase E gene (rpoE, [TIGR00448]), inferred from pre-clustered reference and metagenomic sequences (see Methods). Sequence clusters at tips are annotated according to their source (proGenomes3, GTDB r220 or SPIRE) and the habitat categories from which they were recovered among the full set of metagenomes. Nodes marked with dots indicate clade-level groups at phylum (red) and family (blue) level, inferred based on pre-calibrated cutoffs in relative evolutionary divergence (RED; see Methods). Phylum groups were taxonomically classified based on encompassed sequences from proGenomes3, GTDB r220 or SPIRE MAGs. Nitrosopumilaceae are indicated as an example clade where unbinned contigs substantially enrich known taxa. Deep novel (unclassified) phylum-level clades are highlighted at various points. Phylogeny visualizations for other archaeal marker genes are available under EBI BioStudy S-BSST2111 ^105^.

Marker gene phylogenies recovered previously recognized deep-branching clades as mostly homogenous groups and recapitulated common hypotheses regarding the order of branches near their respective putative roots ^34^. Our estimates of reference phylum-level clades (i.e., containing sequences from proGenomes3 or GTDB) exceeded the number of phyla recognized in the GTDB r220 based on full species trees for both Archaea (estimate: 36±12; GTDB r220: 19) and Bacteria (238±82; GTDB: 175), in part because our RED partitioning algorithm tended to split large clades like the TACK superphylum (represented as single phylum Thermoproteota in the GTDB) or groups predicted to have skewed evolutionary rates such as DPANN archaea into multiple ‘phyla’. At the same time, all explored marker gene phylogenies contained varying numbers of deep (phylum-level) branches that appear novel relative to both proGenomes3 and GTDB r220, i.e. encompassing only sequences in unbinned contigs or SPIRE MAGs. For example, the archaeal rpoE tree in Figure 2 contains, among others, an unclassified phylum-level branch comprising 758 sequences from different habitats, organized into 109 tips (i.e., gene clusters) and five predicted family-level clades, as a sibling group to phylum Thermoplasmatota. Overall, we estimated a median of 10 putative higher-level clades in Archaea (4±3 from SPIRE MAGs, 6±8 among unbinned contigs), corresponding to a ∼28% increase over estimated reference clades, and 145 novel clades in Bacteria (28±29 among MAGs, 117±137 in unbinned contigs), a ∼61% increase over reference estimates. Towards shallower taxonomic levels, our estimates of reference clade counts increasingly approximated those in the GTDB r220 (with the exception of the species level, where only around half of GTDB reference clades were recovered from metagenomic assemblies; see above), whereas estimates on discoverable clades among SPIRE MAGs and unbinned contigs strongly increased with increasing taxonomic resolution (see Figure 3A and Table 1).

**Figure 3.**
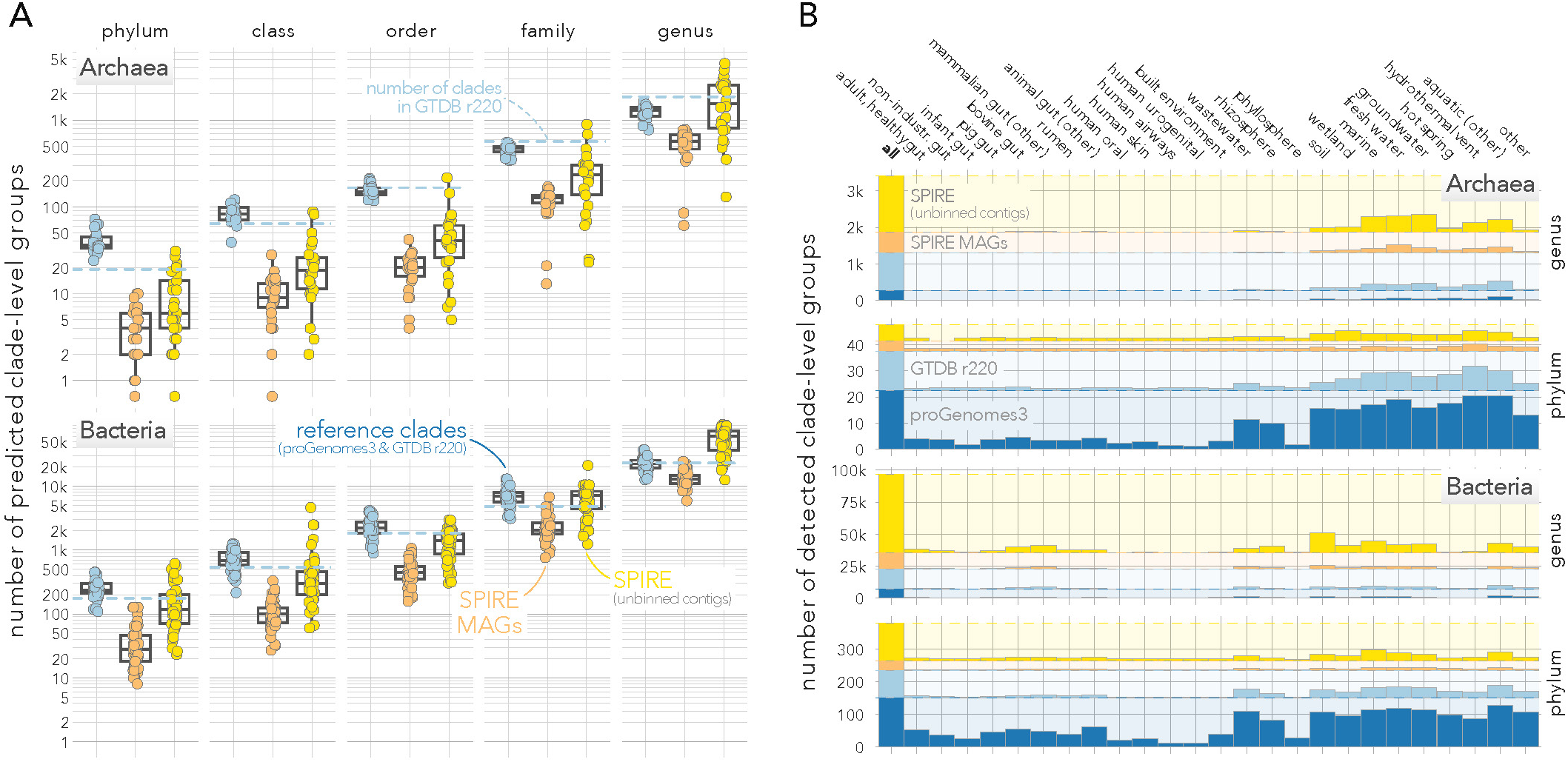
Unbinned contigs and MAGs suggest thousands of discoverable deeper clades. **(A)** Numbers of predicted clade-level groups at phylum, class, order, family and genus level, based on relative evolutionary divergence (RED) cutoffs (see Methods). Each point indicates an estimate for one marker gene. Colours indicate data source: blue for clades containing at least one reference sequence (from proGenomes3 or GTDB r220); orange for those containing no reference sequences, but at least one SPIRE MAG; yellow for clades containing only sequences from unbinned SPIRE contigs. Dotted blue lines indicate the reference number of clades per level in GTDB r220. **(B)** Distribution of discoverable phylum-and genus-level clades across habitats. Bars indicate how many clades for each data source are discoverable across all habitats (leftmost column) or when only considering data from individual habitats.

**Table 1.** Estimated clade counts at deeper taxonomic levels. Clade counts were estimated based on relative evolutionary divergence (RED) cutoffs from individual marker gene phylogenies (see Methods). Values shown are the median ± standard deviation across domain-specific marker gene sets. Note that the discrepancy between reference and estimated species counts is in line with observations that around half of marker gene clusters representing reference species were not observed in metagenomic assemblies (see main text and Extended Figure 3).

The recovery of predicted clades was heterogeneous across habitats (Figure 3B). Gut samples, non-intestinal human sites, and the phyllosphere each covered only a small fraction of both reference and discoverable predicted archaeal and bacterial clades, at both deep (i.e., phylum-level) and shallower (genus-level) resolution. Thus, in spite of comprising a majority of the sequencing effort, these habitats encompassed only a small fraction of known and discoverable diversity. In contrast, the majority of predicted higher-level clades were observable in soils, rhizosphere, wastewater and aquatic environments. Hydrothermal vents stood out as particular hotspots of archaeal diversity: although accounting for just 296 metagenomes in our dataset (0.3% of the total), they contained representatives of four fifths of reference and two thirds of MAG-based and unbinned predicted archaeal phylum-level clades, the widest coverage of deep lineages observed for any tested habitat. Most discoverable genera were predicted in aquatic habitats (marine, freshwater and groundwater) for Archaea and soils for Bacteria. In other words, soils and aquatic habitats remain the major reservoirs of untapped prokaryotic diversity, irrespective of prior sampling efforts.

### Prokaryotic diversity follows Willis’ law and Yule curves across scales of granularity

In 1922, Willis & Yule observed that taxonomic clade size distributions of several plant and animal groups follow power laws: the frequency of genera (i.e., number of species in that genus) decreases with the exponent of genus size ^30^, resulting in *hollow curves* dominated by singleton clades (i.e., genera containing just one species) with a heavy tail of very large clades (i.e., disproportionally species-rich genera). Although adherence to this empirical “Willis’ law” has since been observed for many groups of organisms ^35,36^, is has been suggested that the effect is an artefact of undersampling (rare species are less likely to be ‘discovered’) and biased taxonomic clade definitions (clade boundaries are subjective and anthropocentric ^35,37^). Nevertheless, beginning with Yule’s seminal work postulating preferential attachment of novel species to existing genera, proportionally to genus size ^31^, researchers have sought to identify the evolutionary mechanisms and processes that may give rise to the observable extant biodiversity ^36,38–41^.

We hypothesized that prokaryotic diversity follows the same empirical laws, i.e. that common patterns in biodiversity are shared across all domains of life. We reasoned that our dataset was uniquely suited to test this, as our clade-level groups were data-driven (inferred automatically based on RED values, independently of cultivation and genome binning), taxonomy-agnostic (defined directly from marker gene phylogenies) and comprehensive (sampled across Earth’s habitats at scale). We found that archaeal and bacterial diversity indeed follow Willis’ law: clade frequencies decrease with increasing clade sizes in a power law relationship (Figure 4). Remarkably, this held along the full tested range of >5 orders of magnitude of clade counts – in other words, the frequency distribution was consistent from the largest bacterial clades containing tens of thousands of species to singleton clades with just one species. As shown for the bacterial rsmD gene (16S rRNA guanine methyltransferase, [TIGR00095]) as a representative example, Willis’ law moreover held across taxonomic scales, from species-within-genera to species-within-phyla (Figure 4A) and also for subclades at higher taxonomic levels, i.e. for genera-within-families to classes-within-phyla (Figure 4B). Fitting naïve power law equations for each individual marker gene, we computed a marker gene-specific ‘Willis coefficient ω’ to describe the strength of the relationship (as the slope in a log-log plot; see Methods). We observed a clear trend of decreasing ω (implying a heavier tail, i.e. a stronger bias towards very large clades) across taxonomic levels for species counts (from ω≈1.4 for species-within-genera to ω≈0.1 for species-within-phyla), and more similar ω values in a 1.4-1.7 range for subclade-within-clade counts (Extended Figure 6A).

**Figure 4.**
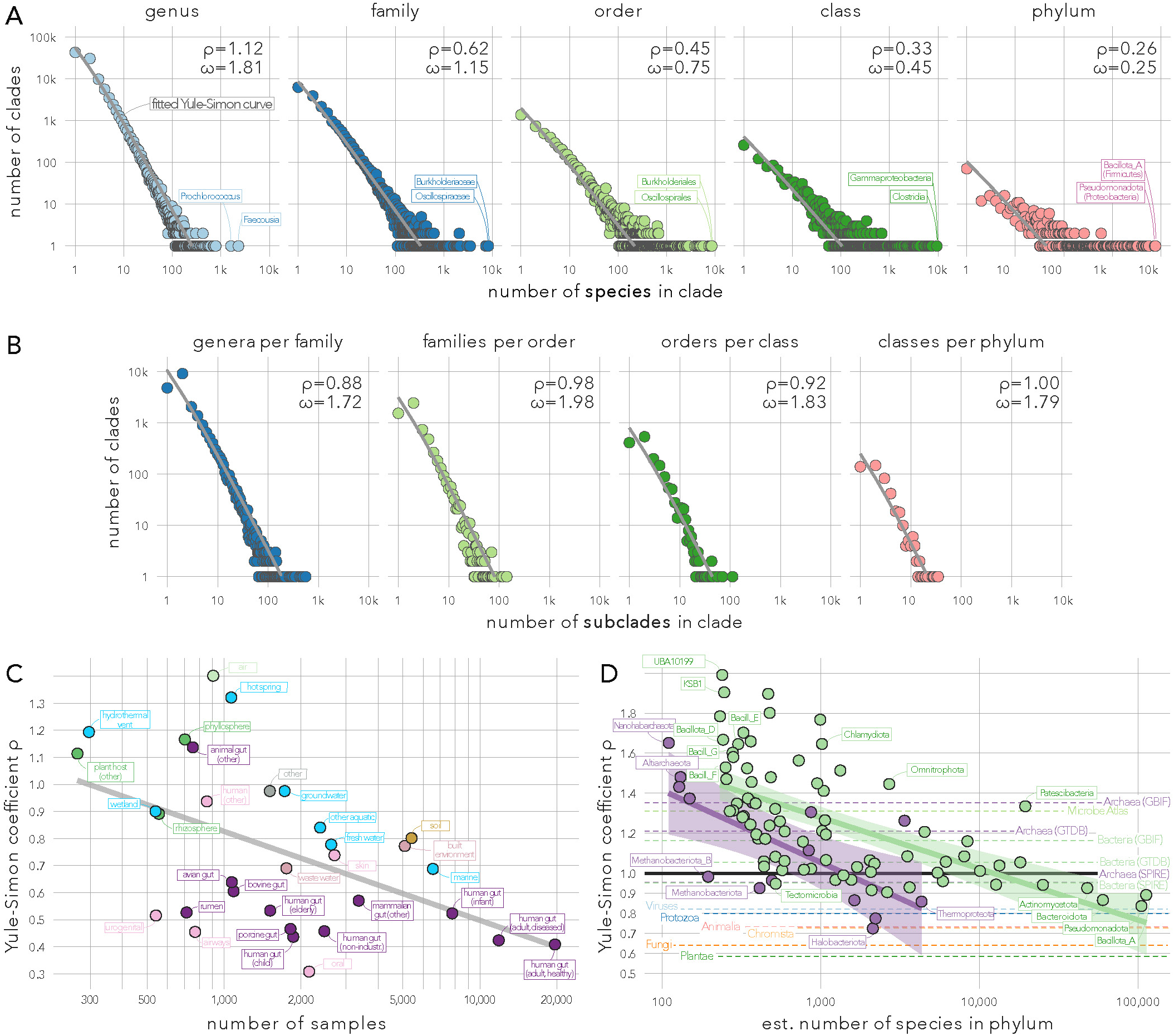
Prokaryotic diversity follows Willis’ law and Yule curves. **(A)** Log-log plot of clade size (number of species per clade; x axis) against clade count (number of clades containing x species; y axis) for the example bacterial marker gene rsmD (16S rRNA guanine methyltransferase, [TIGR00095]). Clade-level groups were inferred based on relative evolutionary divergence (RED) cutoffs from a full phylogeny at each taxonomic level (see Methods). Grey lines indicate fit Yule-Simon curves; estimated Willis coefficient ω and Yule-Simon coefficient ρ are provided (see Methods). **(B)** Equivalent to (A), but showing the number of clades (y axis) containing x subclades (x axis) of the subordinate taxonomic level, i.e. genera-within-families to classes-within-phylum. **(C)** Yule-Simon coefficients estimated for individual habitats, i.e. only considering genes assembled from metagenomes of a focal habitat. X axis corresponds to the number of samples per habitat. **(D)** Yule-Simon coefficients estimated within individual recognized phyla (see Methods). Dotted lines indicate ρ estimated for prokaryotic domains, eukaryotic kingdoms and Viruses based on reference taxonomies from the Global Biodiversity Information Facility (GBIF), the GTDB r226 and the Microbe Atlas Project 16S rRNA OTUs (see Methods). Corresponding Yule curves, analogous to panel A, are shown in Extended Figure 7.

While ‘naïve’ power laws provided good overall fits, we observed that clade size distributions were disproportionately heavy-tailed on the right (i.e., large clades were over-represented) while also slightly deviating from power laws towards the left (i.e., lower clade frequencies for small clade sizes), thereby showing several of the characteristics originally used by Yule to deduce potential underlying processes ^31^. We therefore fit the data to Yule-Simon distributions (see Methods; Figure 4AB). Estimates of the Yule-Simon parameter ρ were mostly consistent across marker genes and between Bacteria and Archaea (Extended Figure 6B): ρ decreased with increasing taxonomic depth for species counts (ρ≈1 for genera to ρ≈0.4 for phyla), with an inverse trend for subclade counts (ρ≈0.85 for genera-within-families to ρ≈1.5 for classes-within-phyla). ρ can be thought of as a ‘rich get richer’ coefficient as it describes the strength of preferential attachment in a Yule process: low ρ values indicate a strong preferential attachment of newly arising subclades to large existing clades. In other words, our results indicate that new species tend to arise in larger existing clades, with a skew towards big clades that is strongest in phyla and least pronounced towards large genera, i.e. the dominance of species-rich phyla (like Bacillota or Pseudomonadota) is more pronounced than that of species-rich genera (like Prochlorococcus). The inverse was observed for subclades within clades, albeit with a less pronounced trend: new genera tend to arise with a stronger bias towards existing large families than new classes towards existing large phyla. Interestingly, for orders-within-classes and classes-within-phyla, we observed a disparity between Archaea and Bacteria (with higher ρ values among Archaea) that is only in part attributable to sampling noise (fewer archaeal clades leading to noisier fits). This may indicate that an archaeal ‘order’ or ‘class’ is not fully equivalent to bacterial ‘orders’ or ‘classes’ in terms of phylogenetic depth, or that indeed archaeal and bacterial diversity at these levels is organised differently.

### Preferential attachment patterns vary across habitats and phylogeny

We explored how clade size distributions are further shaped by environment and phylogeny by refitting curves for individual habitats (Figure 4C) and phyla (Figure 4D). Yule-Simon coefficients were generally lower in well-sampled habitats (ρ_Spearman_ = −0.50 for Bacteria and −0.25 for Archaea), but with clear differences between broader environments (Figure 4C). Intestinal and human non-intestinal habitats showed lowest ρ coefficients, arguably because they are narrowly defined and correspond to just one or few host species. In contrast, higher coefficients were observed for aquatic environments (in particular hot springs and groundwater) and soils, but also non-mammalian guts, indicating that diversity in these habitats is less dominated by few large clades, and that newly discovered species in these environments are more likely to belong to novel deeper clades, whereas they are expected to belong to (known) large clades in the intestine.

Yule-Simon coefficients were strongly associated with total sampled species richness within recognized phyla for both Bacteria (ρ_Spearman_ = −0.71) and Archaea (ρ_Spearman_ = −0.79; Figure 4D): well-sampled phyla like Pseudomonadota or Bacillota_A (formerly Firmicutes) had low ρ coefficients, whereas smaller phyla were less dominated by large clades. Two notable exceptions were Methanobacteriota/Methanobacteriota_B which deviated towards lower ρ (i.e., fewer species sampled, but highly concentrated) and Patescibacteria with an estimated ∼19.5k species distributed into more evenly sized clades (ρ=1.33). Yet nearly all tested prokaryotic phyla had higher Yule-Simon coefficients than eukaryotic kingdoms and even Viruses, inferred based on reference taxonomies of the Global Biodiversity Information Facility (GBIF; see Methods and Extended Figure 7). On the other hand, domain-wide ρ estimates based on our dataset were substantially lower (and more consistent) than those based on GBIF bacterial and archaeal taxonomies, GTDB r226 reference taxonomies ^42^ or on 16S rRNA OTUs from the Microbe Atlas Project reference ^11,43^. This suggests that while eukaryotic diversity (in particular among plants and fungi) is much more concentrated into a few large clades than for Bacteria and Archaea, the organisation of biodiversity follows similar and consistent patterns across the entire Tree of Life.

## Discussion

There are multiple layers of ‘discovery’ when it comes to the description of novel prokaryotic lineages: the isolation of organisms for experimental access and formal taxonomic approval remains standard ^44–47^, yet the delineation of genomes directly from metagenomic sequencing data has enabled the functional and evolutionary characterisation for a greatly extended range of clades without cultivated representatives, and even variants in (amplified) marker gene sequences can be ecologically informative and indicative of further diversity. In this context, our study addresses four basic questions: (i) how many prokaryotic lineages can be discovered in currently available data regardless of limitations of current tools; (ii) how much of this diversity is missed by current genome-centric approaches; (iii) should we expect the discovery of novelty to slow down as more data comes in; and (iv) how is prokaryotic diversity organised across habitats and phylogeny?

Our results extend previous observations ^18,20,23,48^ indicating that isolates significantly underestimate tractable genomic diversity in metagenomes. Yet strikingly, this discovery gap goes both ways: only around half of the species clusters encompassed by genomes of cultivated representatives (and by extension, reference MAGs) in proGenomes3 and GTDB r220 were detected in our metagenomic dataset. These levels of isolate-derived species detection among metagenomes are higher than previous reports ^27^, likely because the present dataset is more comprehensive, although saturating discovery curves for several environments indicate that further detections are not expected when adding more samples (at least when assuming consistent bias in sampling). This tallies with the idea that cultivation efforts and metagenomic sampling often target different (micro-)environments and that cultivability does not necessarily reflect abundance (or ecological importance) in natural communities ^49^, as has been observed for well-studied organisms such as *E. coli* ^50^ or *C. difficile* ^51^.

The vast majority of discoverable diversity in our dataset extended beyond *all* genome-centric approaches. As a lower bound, defined via the independent detection of at least two closely related marker gene sequences in distinct samples (see Supplementary Discussion), one species remains ‘unbinned’ for every species discovered among cultivated organisms (proGenomes3), reference MAGs (GTDB r220) or dataset-specific MAGs (SPIRE). When considering all detected sequence variants, estimates go up to four discoverable species per genomically defined species, or 580k ‘novel’ bacterial and 21k archaeal species, exceeding recent reports of 83k species based on raw read mapping to conserved marker gene windows using SingleM and sandpiper ^28^ or 135k species based on metagenomic assembly ^27^.

At broader taxonomic levels, our data likewise suggests a wealth of discoverable clades in available data: we estimate that less than one in four bacterial genera, and just three in five bacterial phyla that can be inferred from assemblies are currently recognized in reference databases. These are estimates across many phylogenies so that issues affecting individual gene trees (such as paralogs or uneven evolutionary rates, ^52–54)^ are expected to mostly even out. Nevertheless, we note that the proper characterization of novel lineages requires genomic context, or at the very least phylogenies of multiple concatenated informative markers combined with complex models of evolution ^55,56^, beyond the scope of the present study. While these numbers carry increasing uncertainties at broader taxonomic levels, this puts the total (currently discoverable) bacterial phyla in the range of 300-400, compared to 30-40 among Archaea, although we note that RED-based clade definitions at phylum and class levels were broader in Archaea than in Bacteria. Indeed, this ratio of roughly one order of magnitude difference between bacterial and archaeal clade counts held across all taxonomic levels, although towards shallower levels (genus and species), bacterial diversity increased disproportionately. Therefore, our data provides the broadest support yet for the observation that Archaea are significantly less diverse than Bacteria ^7,57,58^. This aligns with the hypothesis that the last archaeal common ancestor, recently estimated to be slightly younger than the bacterial common ancestor ^59^, was a hyperthermophile with the adaptation of archaea to moderate environments suggested to have been supported by gene import from Bacteria ^60^.

At the same time, we found that bacterial and archaeal diversity are organised along common fundamental patterns. Empirical scaling laws are prevalent in microbiology, from biogeographic taxa-area relationships ^61,62^ to macroecological patterns ^63^. Yet to our knowledge, our work provides the first comprehensive evidence that prokaryotic clade size distributions follow power laws, supporting century-old hypotheses by Willis ^30^ and Yule ^31^. Power laws are a hallmark of fractal geometries, and the remarkable consistency of inferred coefficients not only between Bacteria and Archaea, but also across eukaryotic kingdoms and viruses and even for subclades-within-clades at different depths imply that biodiversity is indeed fundamentally fractal: that consistent (and possibly predictable) evolutionary forces mould the Tree of Life into recurring patterns from root(s) to tips ^38,64–66^. Our data strongly support Yule’s hypothesis ^31,36^ that one central pattern is preferential attachment: ‘the rich get richer’ as new clades arise preferentially within already large clades. The effect varies across habitats and phylogeny and can also be interpreted in light of the discovery of novel clades: assuming random sampling, far more new species need to be ‘discovered’ in intestinal environments or species-rich phyla like Pseudomonadota (with low Yule-Simon coefficients) than in hot springs or among Patescibacteria (higher coefficients) to identify a novel deeper clade. From a purely technical standpoint, we moreover note that our observations provide strong support for the use of relative evolutionary divergence (RED) to delineate archaeal and bacterial clades ^32^, as our mostly agnostically inferred RED-based clades fall into the described consistent patterns.

We note that although our methodology includes filtering and correction for various error sources, and although assembled sequences greatly underestimate the total sequence variation represented among raw metagenomic reads ^67^, the presence of assembly and clustering artefacts and spurious gene variants (including paralogs, or eukaryotic or viral orthologs) are possible sources of noise that may lead to an overestimation of diversity. Either way, the gap between genomically captured and unbinned discoverable diversity was growing (rather than closing) in virtually all tested habitats. And given that most metagenomically derived genes ^67^ and inferred gene families ^68,69^ are likewise unaccounted for by genomes, our results define a range of how many additional lineages may be genomically disentangled from the vast space of unbinned and unclassified reads with improved algorithms. Indeed, constantly refined sampling and sample processing protocols, the increasing use of longread technologies ^70,71^, faster mapping tools that render multi-sample co-binning more computationally tractable ^72,73^, a novel generation of binning tools ^74–76^ and innovations like the iterative targeted co-assembly workflow in Bin Chicken ^29^ promise to make major inroads into this bulk of untapped diversity.

Our dataset reflects existing sampling bias both *between* habitats (human fecal samples dominate the survey) and *within* environments (uneven coverage geographically, along depth-elevation gradients and with regard to host species) as individual datasets are usually generated for specific purposes, rather than as part of global surveys. Indeed, the habitat definitions used in the present work are – by design – pragmatic and guided by data availability, and not necessarily reflective of the underlying diversity of environmental conditions ^77^: for example, soils are structured into many heterogeneous sub-environments with physicochemical and biotic parameters that vary at micrometer scale ^78,79^, whereas well-mixed ocean layers can present homogeneous environments across several square kilometers ^80^. Moreover, while we use metagenomic samples as units of reference, we note that established protocols capture very different scopes of underlying communities: while typical ocean water samples (filtered from 10-100L ^80^) and human fecal samples (∼1g of feces) carry total microbial loads of 10^10-10^11 cells ^81^, typical soil samples of 1-2g ^82^ contain 2-3 orders of magnitude fewer cells ^2^. Given that Earth is expected to harbour a total of ∼10^29 microbial cells each in ocean water and soils, but just 10^23-10^24 cells in the global human gut microbiome ^2^, this puts both the abscissa (i.e., the number of samples considered) and the ordinate (i.e., the number of clades discovered) in the presented rarefaction curves into further perspective. Finally, both physiological and technical factors may reduce (or bias) recall: among others, different membrane and cell wall types yield differentially to common extraction protocols ^83^, genomic composition (including GC content) and structure impact sequencing protocols ^84^, and current gene calling algorithms still often struggle with non-standard genetic codes ^69^.

We therefore caution against extrapolating total species richness estimates from our rarefaction curves, as future data may break out from previous sampling bias by adding samples from uncharacterized sub-environments. This also means that our inferred rarefaction curves and novel species discovery coefficients should be considered conservative lower bound estimates, as they assume continued sampling from the same underlying distribution, while in fact the vast majority of the planet remains metagenomic *terra incognita* and overlooked endemic clades in unsampled sites are expected to disproportionately contribute to global biodiversity ^85,86^. In spite of these caveats, soils, aquatic habitats and non-mammalian guts clearly stood out among studied habitats as both key reservoirs of discoverable diversity in existing data and as hotspots primed for future discovery, and our data suggest that for almost all considered habitats, the discovery of novel diversity will remain in full swing for the foreseeable future.

The present work takes stock of characterised and discoverable microbial diversity based on a snapshot of available data. It is encouraging – and, from a data science perspective, slightly intimidating – that the underlying data continues to accrue near-exponentially in public repositories. But as we have shown, even existing data may hold countless surprises yet, as a vast amount of discoverable prokaryotic diversity is ‘hiding in plain sight’, flying under the radar of current genome-centric approaches. Most undiscovered diversity – both that within existing data and that not yet sampled – will likely be part of the ‘rare biosphere’, and thus cover a disproportionately large functional diversity (of potential medical and biotechnological relevance), but will also follow predictable global patterns of diversity across habitats and phylogeny that can suggest where sampling effort should be directed. Cataloguing and characterizing these organisms, with a focus on understudied environments, is therefore not only relevant to the study of microbial and ecology and evolution, but can be considered an integral aspect of global conservation efforts.

## Methods

### Data sources and preprocessing

We used three principal data sources in our analysis: (i) proGenomes v3 ^16^, a quality-filtered subset of 907,388 isolate genomes sourced from NCBI’s RefSeq and GenBank databases ^87^ organised into 41,171 species-level clusters based on a set of 40 ‘specI’ marker genes ^12^; (ii) the Genome Taxonomy Database (GTDB) release 220 (April 2024, ^17^), a curated set of 596,859 isolate and metagenome-assembled genomes (MAGs), clustered into 107,235 bacterial and 5,869 archaeal species-level clusters sharing ≥95% whole-genome Average Nucleotide Identity (ANI); and (iii) SPIRE v1.1 ^23^, encompassing a curated set of 92,187 shotgun metagenomic samples assembled into ∼23.3Tbp of contigs, a subset of which were binned into 1,158,468 medium or high quality MAGs, organised into 107,078 species-level clusters that partly overlap with those in proGenomes3.

Genomes from all three datasets were downloaded and taxonomically (re-)classified against the GTDB r220 using GTDB-tk v2.4.0 ^88^, and consensus taxonomies for species-level clusters were inferred based on adjusted majority votes, as described previously ^23^. Compared to SPIRE v1.0, 6,959 metagenomic samples were excluded for the present analysis based on data type and provenance (e.g., excluding samples explicitly enriched for viruses) or insufficient assembly size. For the remaining 92,187 samples, habitat annotations and contextual data were updated by manually curating against an extended *microntology* v0.3.0 encompassing 103 terms and categories (reference set and updated annotations available via https://spire.embl.de/downloads and table S2) and further organised into 32 higher-level categories (see table S1).

### Estimates of bacterial and archaeal diversity based on taxonomic marker genes

The majority of analyses presented in the main text are based on 168 near-universal taxonomic marker genes as established by the GTDB: 120 bacterial (‘bac120’, ^13^) and 53 archaeal (‘arc53’, ^14^) markers, with an overlap of five genes. We downloaded profile Hidden Markov Models (HMMs) for these marker sets as part of the GTDB-tk v2.4.0 database and used the HMMER v3.4 ^89^ *hmmsearch* routine to identify and extract marker gene sequences among predicted Open Reading Frames (ORFs). For the final filtered set of marker genes (see below), this yielded on average 904,654±98,879 sequences per bacterial marker in proGenomes3, 93,198±4,733 among GTDB r220 species representatives, 840,023±133,904 in medium and high-quality SPIRE MAGs, and 3,051,465±1,290,024 among ‘unbinned’ SPIRE contigs, i.e. those not assigned to a MAG passing genome quality filters in SPIRE v1. For Archaea, we obtained 2,451±150 sequences from proGenomes3, 4,898±410 from GTDB r220 species representatives, 15,504±2,593 from SPIRE MAGs and 107,419±75,823 from unbinned SPIRE contigs (see table S3 for a per-gene overview).

For each marker gene, we clustered extracted sequences using the MMseqs2 cascading clustering routine ^90^ with a cutoff of 96.5% sequence similarity (see below) and otherwise default parameters. To convert the number of marker gene sequence clusters into a corresponding number of species-level clusters, we estimated conversion factors as follows. First, we generated marker gene cluster discovery curves via iterative logarithmic rarefaction, i.e. we downsampled the number of considered gene sequences along a logarithmic scale (10, 20, …, 100 sequences; 200, 300, …, 1000 sequences; etc) with 10 iterations at each step. At each rarefaction point, we recorded the number of ‘discovered’ marker gene sequence clusters and the number of represented species or species-level genome clusters in proGenomes3, GTDB r220 and among SPIRE MAGs, considering each data source individually. We then used linear regression models of the type *number_of_species ∼ number_of_gene_clusters* with a forced intercept at 0 along these rarefactions to estimate gene cluster to species conversion factors (i.e., the number of newly discovered species per newly discovered marker gene cluster). Extended Figure 8 shows the resulting fits for 10 randomly selected marker genes each for Archaea and Bacteria. Based on benchmarks of marker gene sequence similarity cutoffs ranging from 95% to 99.5%, we based further analyses on 96.5% clusters as these showed very robust linear fits (with standard errors in the range of 10^-3) with conversion factors closest to identity (i.e., roughly one species discovered per marker gene cluster discovered), and high consistency across the different species-level reference clusterings in the underlying datasets (based on 40 specI marker genes in proGenomes3, 95% whole-genome ANI in GTDB and a combination of both approaches among SPIRE MAGs). Finally, we filtered the set of considered marker genes by excluding redundant markers (part of both the arc53 and bac120 sets and therefore prone to cross-mappings) and by removing markers for which predicted species numbers deviated by more than 20% from the domain-level median (in particular for some archaeal markers, detected sequence counts were inflated, possibly due to the detection of bacterial paralogs). Sequence counts per data source for the remaining set of 28 archaeal and 102 bacterial marker genes are available in table S3.

### Inference of global and habitat-resolved species discovery curves

For each of 32 broadly defined habitat categories (ranging from 267 to 19,659 samples per group, see table S1), as well as for the combined set of all 92,187 metagenomes in SPIRE v1.1, we generated bacterial and archaeal species discovery (or rarefaction) curves. We iteratively subsampled the number of considered metagenomes along a logarithmic rarefaction scale, with 5 random permutations per step. At each rarefaction point and for each considered marker gene, we inferred the number of discovered species based on the number of discovered marker gene clusters, using gene-specific conversion factors as described above. Rarefaction permutations were averaged per marker gene and then summarised per marker gene set (bac120 for Bacteria and arc53 for Archaea) as median predicted species counts at each rarefaction step across marker genes (see Figure 1). Extended Figure 1 shows relative errors (coefficient of variation, i.e. standard deviation divided by mean) at each rarefaction step for each marker gene; consistently, the coefficient of variation decreased to <5% when including 1k-10k samples for Bacteria (i.e., when using ∼1-10% of total data), with drops at slightly higher rarefaction steps for Archaea.

In an independent approach, we performed an ‘incremental’ rarefaction, stratified by habitats: we sequentially added samples from each of the 32 broadly defined habitat categories, starting with ‘adult, healthy human gut’ samples, randomized discovery order within each habitat block and then tracked the *globally* discovered species (see Figure 2 and Supplementary Figures 1 and 2). In other words, we tracked the contribution of each habitat to overall discovered species diversity, beyond the diversity discovered in previously considered habitats.

Counts of discovered marker gene clusters (and, by proxy, inferred discovered species) were hierarchically stratified by data source to account for overlap between datasets as follows: all clusters containing at least one sequence originating from a genome in proGenomes3 were labelled as ‘proGenomes3’, irrespective of the origin of other sequences within the same cluster (data series marked in dark blue in main figures 1 & 2); clusters containing sequences from the GTDB r220 (and any other data source *except* proGenomes3) were labelled as ‘GTDB’ (light blue in figures 1 & 2); clusters containing sequences from SPIRE MAGs (but neither from proGenomes3 nor GTDB) were labelled as ‘SPIRE MAG’ (orange data series); clusters containing only sequences from unbinned SPIRE contigs were labelled as ‘unbinned, non-singleton’ if they contained at least two sequences from different contigs (dark yellow data series), and ‘unbinned, all’ otherwise (light yellow data series). Thus, clusters labelled as ‘GTDB’ represent sequence diversity contained in GTDB r220 *beyond* that already represented in proGenomes3, while a significant subset of ‘proGenomes3’-labelled clusters also contained sequences originating from GTDB, SPIRE MAGs or unbinned contigs. Similarly, ‘SPIRE MAG’ clusters corresponded to diversity *not* already covered by proGenomes3 and GTDB r220; and ‘unbinned’ clusters encompassed diversity not represented in any genome or MAG in the dataset. Moreover, by design we only tracked marker gene clusters in metagenomic samples, meaning that a non-negligible subset of genome-based clusters from proGenomes3 and GTDB r220 were not represented in rarefaction runs and calculations, as corresponding sequences were not detected in any of the 92,187 considered metagenomic assemblies (see main text).

### Calculation of species discovery coefficients

We quantified the rate at which novel species-level clusters are discovered in each habitat (and globally, across all habitats) using the equation

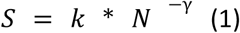

where S is the number of newly ‘discovered’ species per N samples added to the survey; k is a proportionality constant; and γ is a saturation coefficient. An analogous formula is commonly used to describe a (microbial) pangenome’s *openness* ^91,92^, i.e. the degree to which novel genes are discovered as more genomes from the same species are considered. Moreover, the approach is conceptually and mathematically related to the Heridan-Heaps law in linguistics which describes the number of distinct words in a document as a function of that document’s length ^93^.

We fit equation (1) to each habitat-specific and global rarefaction curve described above, stratified by data source, resulting in estimates for γ for each considered marker gene; Extended Figure 9 shows the example fits for a randomly selected subset of marker genes. For more intuitive interpretability, we calculated a *species discovery coefficient* α as

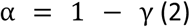

and summarised values across marker genes within each domain-specific set as median values. Thus defined, α scales on [-∞, 1], although only mildly negative values are expected to be observed in practice. If α ≤ 0, species discovery in a habitat is *saturated*, meaning that adding more samples of the same type is not expected to add novel species to the survey – analogous to a ‘closed’ microbial pangenome where additional strains do not add novel genes. Values for α in [0, 1] correspond to *unsaturated* species discovery curves where additional samples continue to add novel species to the survey (analogous to ‘open’ pangenomes); lower α values indicate a more pronounced ‘flattening off’ in the rarefaction curve, indicating a more pronounced slowdown in novel species discovery; higher α values indicate a less pronounced decrease in the rate of species discovery. For α ➝ 1, species discovery is *fully unsaturated*, meaning that each newly added sample adds novel species to the survey, with no discernible ‘flattening off’ in the species discovery curve.

### Phylogenetic inference and analyses

Phylogenetic trees for each marker gene were constructed as follows. We first re-aligned cluster representative amino acid sequences against the respective marker gene HMMs using HMMER v3.4’s ^89^ *hmmalign* routine, trimmed the resulting pseudo-multiple sequence alignments to informative columns using clipkit ^94^ in ‘kpi-gappy’ mode and removed sequences with >70% gaps among the remaining columns. Based on the resulting alignments, we inferred phylogenetic trees using FastTree2 ^95^ under the Whelan-Goldman model ^96^.

We calculated *relative evolutionary divergence* (RED) values ^32^ for each node in the resulting trees using the castor package v1.8.3 in R ^97^. To infer marker-gene specific relative evolutionary divergence (RED) cutoffs at phylum, class, order, family, and genus levels, we subset phylogenies to sequences originating from fully taxonomically classified genomes in proGenomes3, the GTDB r220 or among SPIRE MAGs, cut the trees at incremental RED values (with a RED tolerance of ±0.1) and compared the resulting partitions (i.e., tip sets) versus assigned taxonomic labels, quantifying partition consistency as *adjusted mutual information* (AMI) ^98^. These steps were performed iteratively on trees re-rooted using each recognized phylum in turn as an outgroup and then summarised across alternative root positions (ignoring the current outgroup) to obtain average RED values per internal node, in a workflow adapted from ^32^.

After confirming that the resulting RED cutoffs at peak AMI (i.e., best capturing the distribution of established taxonomic labels) well approximated those reported in ^32^ across individual archaeal marker genes, we used GTDB’s reference cutoffs (with a ±0.1 tolerance interval) for bacterial marker gene trees and cut the full trees at the respective levels. The number of resulting clusters (i.e., nodes where the tree was cut) and their composition (i.e., each node’s descendant tips) were then used as estimates of clade-level groups at each taxonomic level, for each marker gene. We manually selected genes that showed the most consistent and robust profiles across iterations (and rootings) and whose phylogenies were not ostensibly impacted by paralogs, bringing the final sets of considered genes to 29 archaeal and 53 bacterial markers. Marker gene phylogenies were visualized using the ggtree and ggtreeExtra packages in R ^99^.

Marker gene trees were also used for the analyses underlying cumulative marker gene distributions with results presented in Extended Figure 4. Internal nodes corresponding to phyla recognized in the GTDB were identified as the most recent common ancestors of tips representing marker gene clusters containing genes from reference genomes or MAGs classified to the focal phylum. Phylum-level subtrees cut at the respective MRCA nodes were then profiled for the ratio of tips representing only unbinned genes to tips representing genomes (i.e., ‘binned’ genes), as well as for average genomic features (GC content, genome size, coding density) and the estimated total number of represented species. To alleviate effects of genome misclassification and errors in phylogenetic inference, analyses were repeated 5 times for each phylum, each time using a random subset of 80% of genomes to identify the MRCA node.

### Fitting taxonomic clade size distributions

Four datasets were used to fit clade sizes to previously suggested distributions: (i) the above-defined clade-level groups (based on RED cutoffs), for each archaeal and bacterial marker gene; (ii) the GTDB r226 archaeal and bacterial reference taxonomies (accessed via gtdb.ecogenomic.org); (iii) the Global Biodiversity Information Facility (GBIF) backbone taxonomy for eukaryotes, bacteria, archaea and viruses ^100^, accessed using the taxizedb R package ^101^; (iv) the Microbe Atlas Project ^11^ 16S rRNA reference database v3.0 ^43^, using 97% OTUs to represent ‘species’ and 90% OTUs to represent ‘genus’. For each of these taxonomies, we first fit “Willis’ curves” ^30^ relating the frequency of clades to their size (i.e., the number of subclades they contain) as a regular power law:

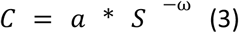

where *C* is the number of higher clades (e.g., genus) containing *S* sub-clades (e.g., species); *a* is a proportionality constant; and ω is the scaling coefficients, referred to as *Willis coefficient* here for simplicity. In other words, (3) relates the frequency of a clade to its size, with positive ω values indicating that large clades (e.g., genera containing many species) are exponentially less common than small clades (e.g., genera containing few or only one species). Otherwise, given the similarity of eq (3) to eq (1) above, the interpretation of ω is closely related to that of the above-defined species discovery coefficient α.

We moreover fit clade sizes to the distribution first suggested by Yule ^31^ specifically for this purpose and later formalized by Simon ^102^:

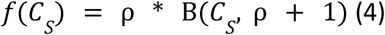

where *f(C_S_)* is the non-negative frequency of clades of size S (i.e., containing S sub-clades); Β is the beta function; and ρ > 0 is the Yule-Simon shape parameter. The distribution results from a Yule process where newly arising subclades preferentially attach to larger existing clades, proportionally to the existing clades’ sizes, sometimes colloquially referred to as a “rich get richer” process. For sufficiently large counts C_S_, the frequency f(C_S_) approximates a power law as in eq (3), with coefficient ω ∼ ρ + 1. Parameter ρ indicates the shape of the distribution and the strength of ‘preferential attachment’, with large ρ indicating a sharper drop in the distribution (less pronounced preferential attachment, distribution less tail-heavy, higher dominance by small clades), whereas smaller ρ indicate more tail-heavy distributions and stronger preferential attachment effects (large clades are larger, small clades are fewer). In other words, ρ can be thought of as a “rich get richer” coefficient, with low ρ values indicating a stronger dominance of fewer large clades in the clade size distribution.

Habitat-stratified analyses (Figure 4C in the main text) were conducted by recomputing clade size distributions using only genes assembled from samples of the focal habitat. For phylum-resolved analyses (Figure 4D), subtrees corresponding to recognized phyla were iteratively extracted from full marker gene phylogenies (see above) and clade size distributions recomputed. In both cases, estimated ρ values were summarised as the median across marker genes.

## Supporting information

Supplementary Discussion

## Acknowledgements

The authors thank Askarbek N Orakov (Harvard T.H. Chan School of Public Health) and Jens Walter (University College Cork) for helpful discussions and feedback on the manuscript. This research was conducted with the financial support of Research Ireland under Grant Number 12/RC/2273-P2 (V.P.P.K and T.S.B.S) and from the Australian Research Council (grant #FT230100724, to L.P.C). A.S. has received funding from the European Research Council (ERC) under the European Union’s Horizon 2020 research and innovation programme (grant agreement No. 947317, ASymbEL), which also supported the position of O.M. This work made use of EMBL IT Services HPC resources ^103^.

## Data Availability

Metagenomic assemblies, Metagenome-Assembled Genomes (MAGs), gene calls and corresponding annotations are available via spire.embl.de/downloads. Extracted marker gene sequences for the ar53 and bac122 sets from SPIRE assemblies, proGenomes3 and GTDB r220 are likewise available via spire.embl.de/downloads. Pre-processed and derived data is available via Zenodo ^104^. Inferred marker gene phylogenies with annotations, as well as pre-generated tree visualizations for archaeal markers are available via the EBI BioStudies repository under accessions S-BSST2111 ^105^, S-BSST2112 ^106^, S-BSST2113 ^107^, S-BSST2116 ^108^, and S-BSST2117 ^109^.

## Code Availability

Supporting analysis code has been deposited under https://github.com/grp-schmidt/ms-census.

## Extended Data Figure Legends

**Extended Data Figure 1.**
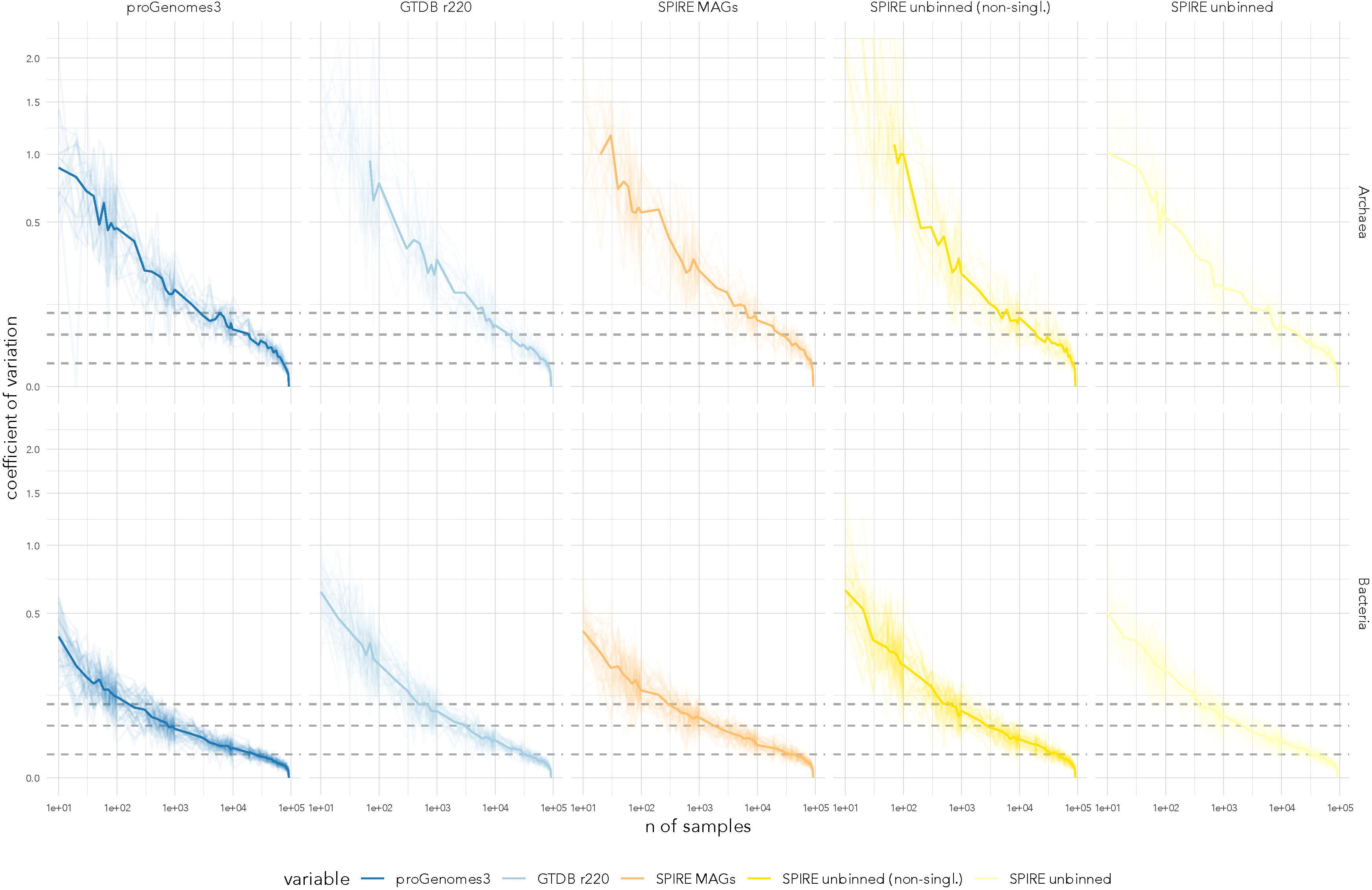
Uncertainty and noise in marker gene-based rarefaction curves. The coefficient of variation (*cv*; standard deviation divided by mean; y axis) across 5 random permutations is shown per rarefaction step (x axis) for the different tested data sources. Thin data series correspond to individual marker genes, the median across marker genes is emphasized. Dotted lines indicate reference levels of 1%, 5% and 10% cv. Note that the sharp drop in cv towards the right is a mathematical necessity as noise between permutations decreases once nearly all samples are included.

**Extended Data Figure 2.**
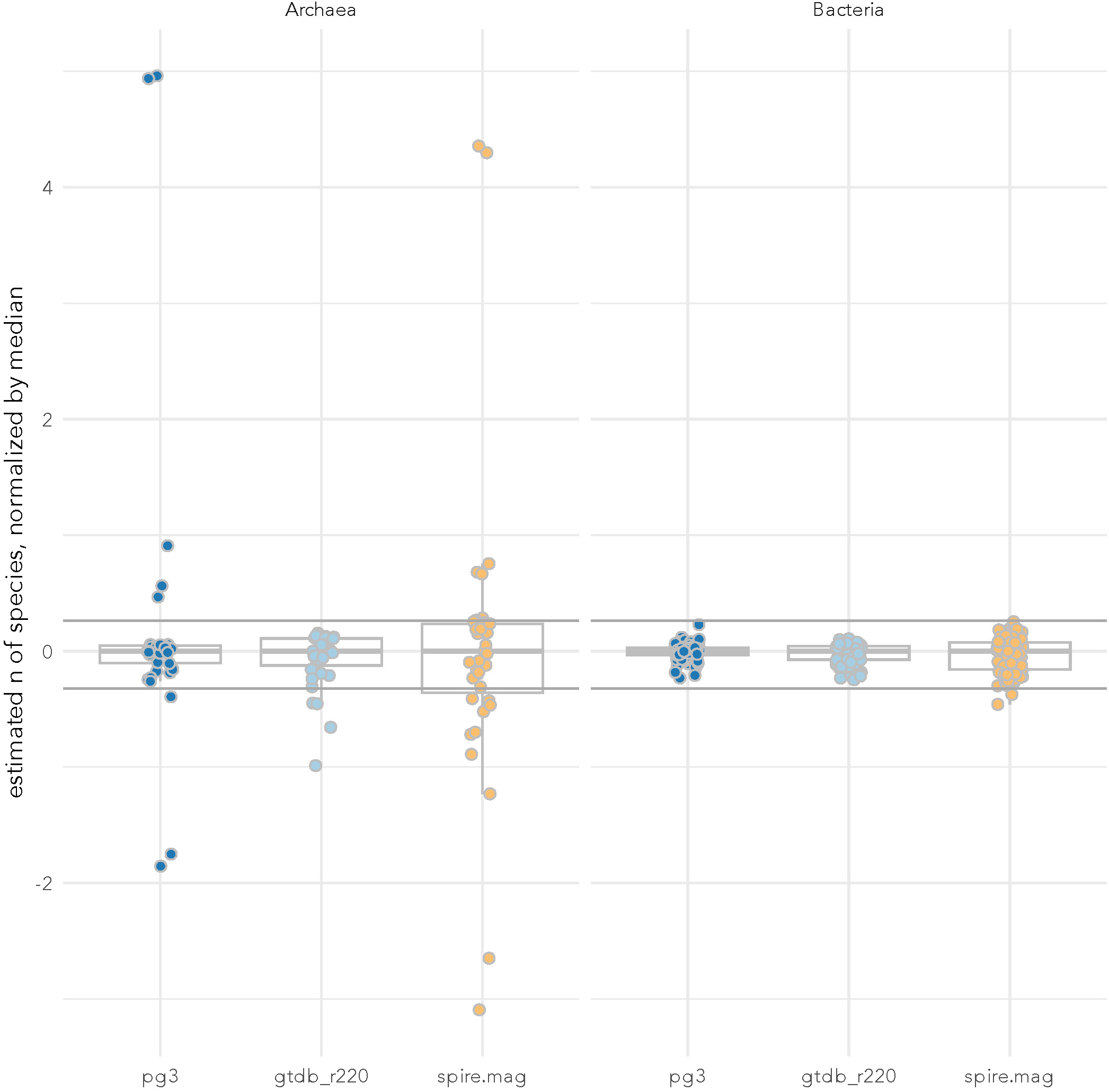
Species count estimates are remarkably consistent across different marker genes. Each dot represents the estimated number of species based on observed marker gene clusters (see Methods) for one marker gene, normalized by the median across marker genes within categories. Archaeal outliers to the top were marker genes with putative cross-mapping to bacterial orthologs (leading to an overestimation of diversity). For analyses in the main text, only a subset of consistent marker genes, further filtered based on additional criteria (see Methods) was used.

**Extended Data Figure 3.**
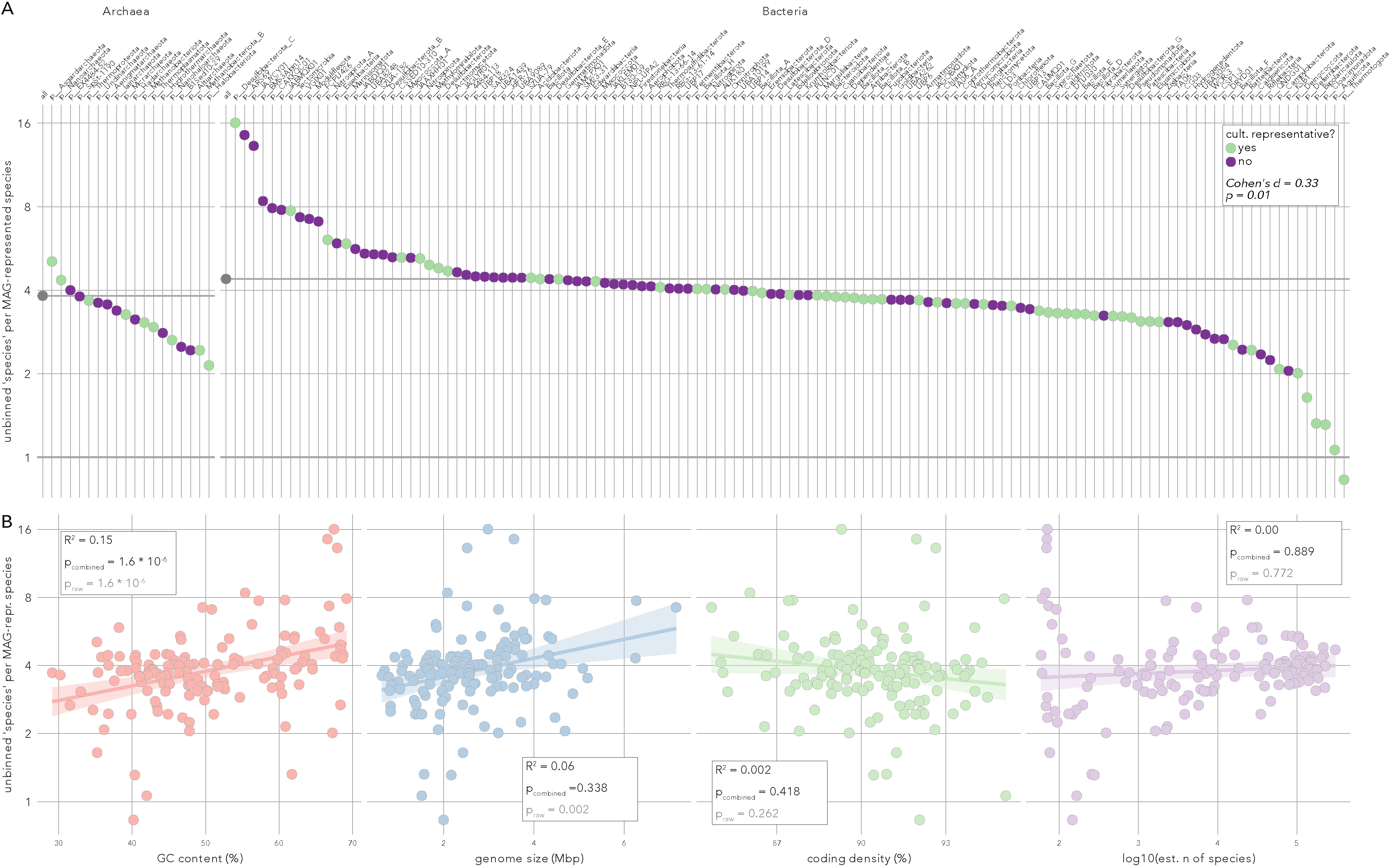
Unbinned genes are enriched among uncultivated phyla and associated with genomic GC content. **(A)** The ratio of tips (i.e., marker gene clusters) containing only unbinned genes to those containing genes from reference genomes or MAGs (y axis) is shown for subtrees of individual recognized phyla, sampled iteratively from full marker gene phylogenies (see Methods). This unbinned ratio was higher for phyla without (purple) compared to those with (green) cultured representatives (Cohen’s d = 0.33; Wilcoxon p = 0.01). **(B)** Ratio of unbinned to binned markers per phylum (y axis) shown against average genomic GC content, genome size, coding density (all derived from GTDB reference genomes per phylum) and the estimated number of species per phylum. R^2^ and p values are shown for individual linear regressions of unbinned ratios against each variable, as well as for a combined multiple regression in which only GC content stood out as significantly associated in an ANOVA.

**Extended Data Figure 4.**
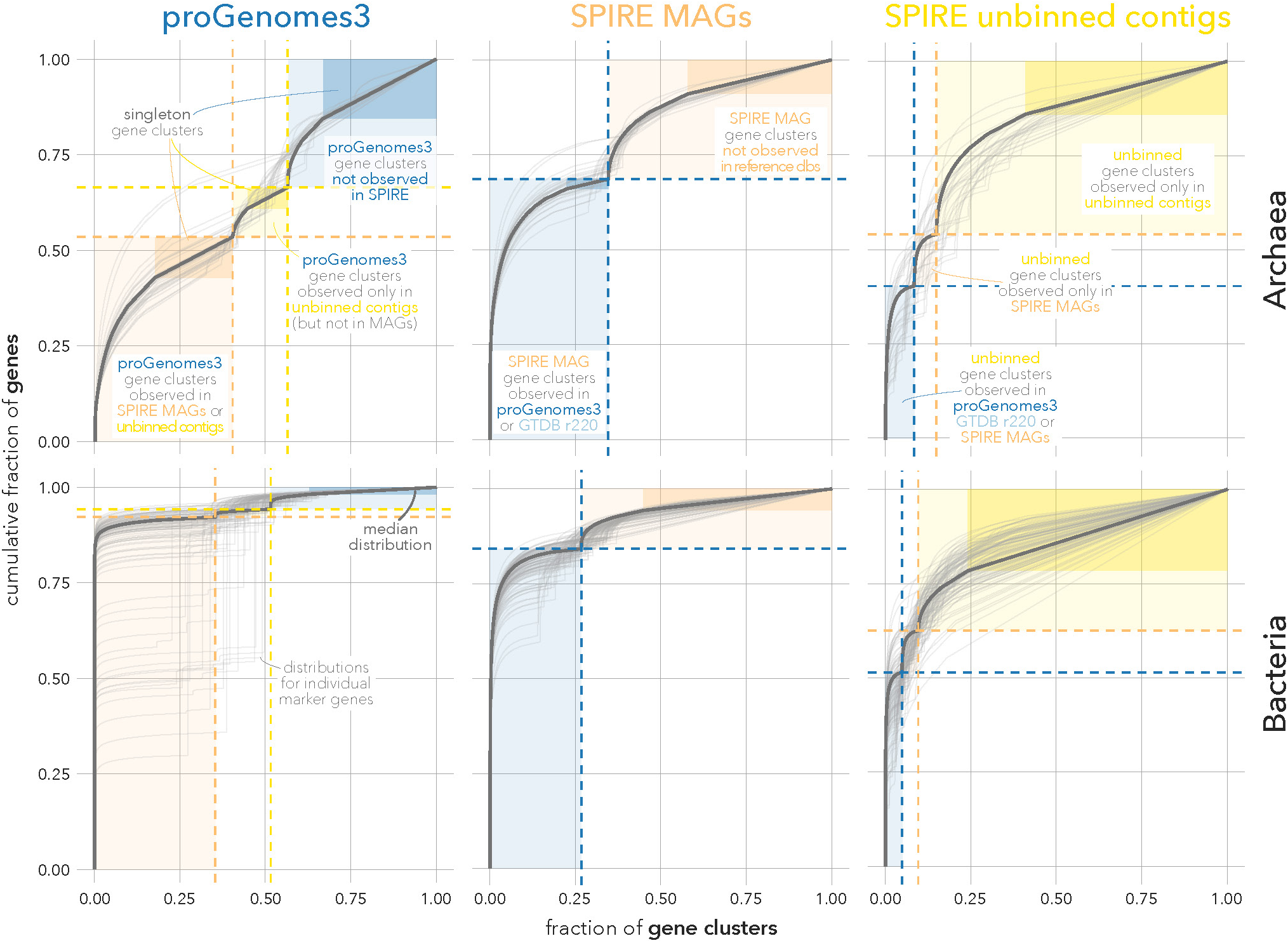
Cumulative distributions of marker genes per marker gene cluster. The cumulative fraction of marker genes (y axis) contained in ranked marker gene clusters (x axis) is shown for individual marker genes (grey lines) and the median marker gene (emphasis). Emphasized rectangles indicate the part of the distribution corresponding to singleton marker gene clusters. The left panels show distributions for marker genes extracted from **proGenomes3** reference genomes: the leftmost part (orange) corresponds to gene clusters also observed in SPIRE MAGs or unbinned contigs; the middle section (yellow) correspond to clusters observed in unbinned contigs, but not MAGs; the rightmost section (blue) are clusters not observed in metagenomic assemblies. The central panels show distributions for marker genes extracted from **SPIRE MAGs**: marker gene clusters that were also observed in proGenomes3 or GTDB r220 genomes (blue, left) or not (right, orange). The rightmost panels show distributions for marker genes extracted from **unbinned contigs**: gene clusters also observed in reference genome sets (blue, left); those observed in SPIRE MAGs, but not reference genome sets (yellow, central); and those observed exclusively in unbinned contigs.

**Extended Data Figure 5.**
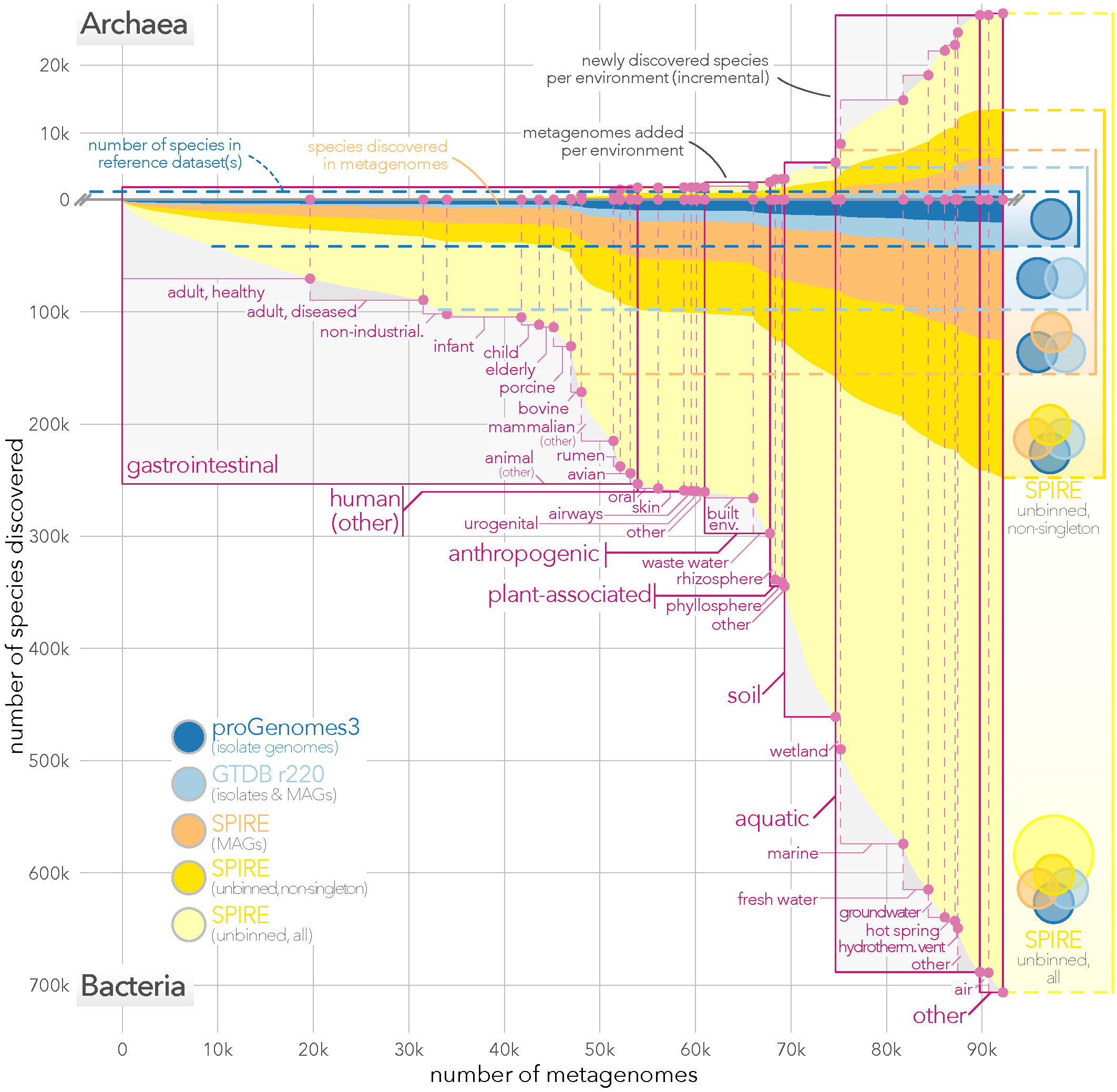
Soils and aquatic habitats are major reservoirs of unexplored microbial diversity. ‘Habitat-cumulative’ rarefaction curves for Archaea (top) and Bacteria (bottom). Samples were added to the survey sequentially by habitat and newly discovered species were tracked *incrementally*, i.e. in addition to those contained in previously added habitats (see Methods). Horizontal dotted lines indicate reference levels, i.e. species counts including those not detectable in our metagenomic dataset.

**Extended Data Figure 6.**
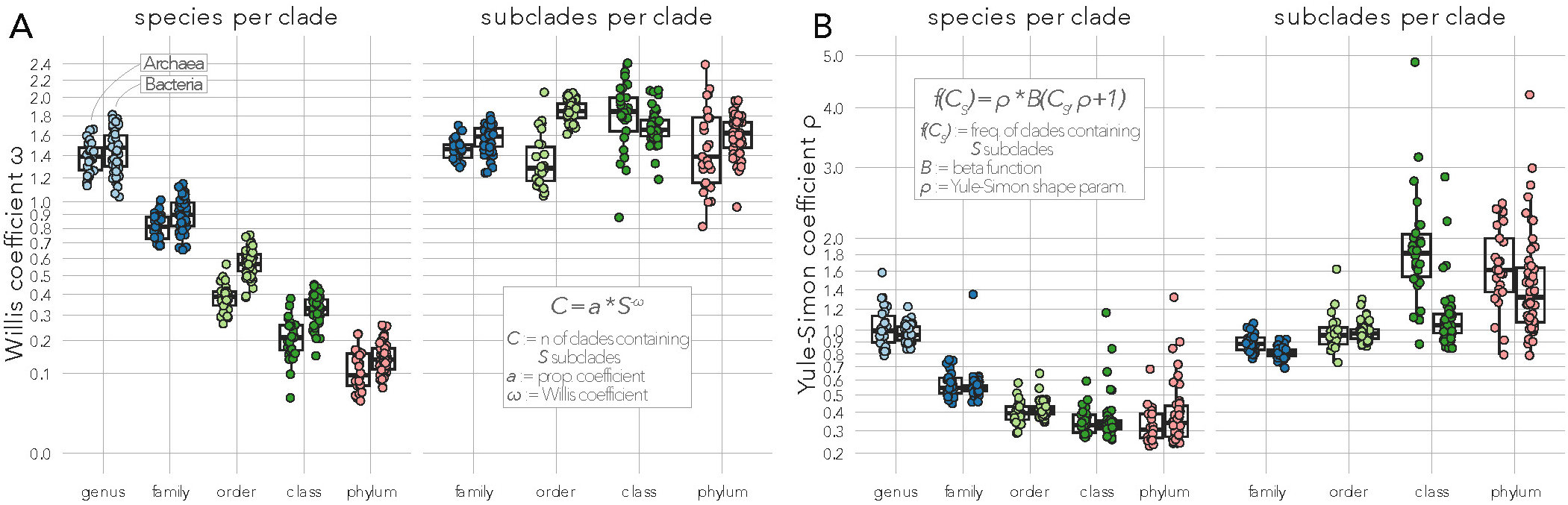
Willis and Yule-Simon coefficients across marker genes and taxonomic levels. Estimated Willis coefficients ω (**A**) and Yule-Simon coefficients ρ (**B**) across taxonomic level. Each dot represents an estimate for one individual archaeal (left series) or bacterial (right series) marker gene. Data is provided for both ‘*species per clade*’ (clade size distributions defined as the total number of species per clade, i.e. per phylum, class, etc) and ‘*subclades per clade*’ (i.e., genera per family, families per order, etc).

**Extended Data Figure 7.**
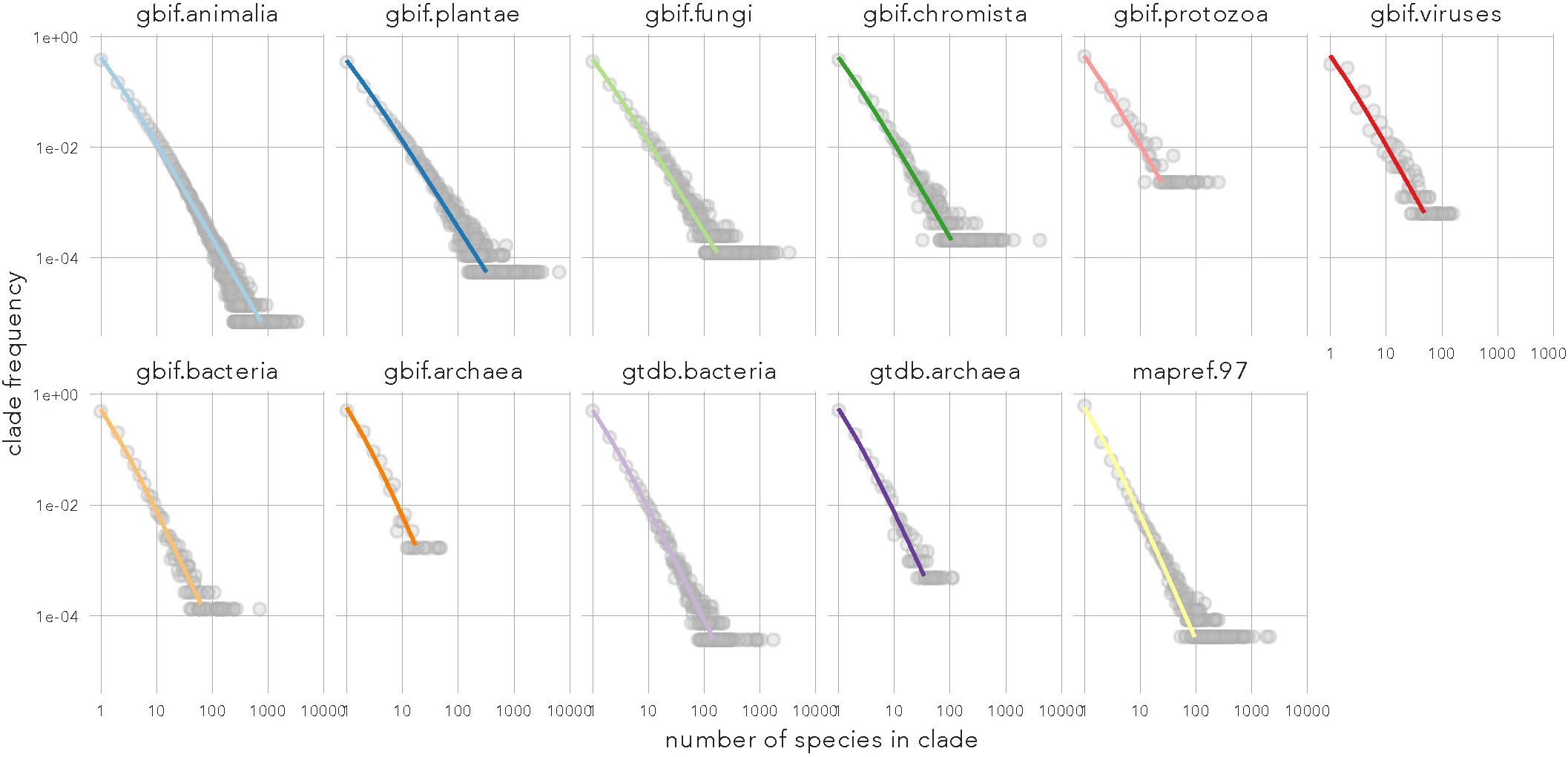
Yule curves based on external reference taxonomies. Log-log plots of clade sizes (number of species per clade; x axis) against clade counts (number of clades containing x species; y axis), analogous to Figure 4A. Lines correspond to fitted Yule curves. Each data series corresponds to a reference taxonomy from the Global Biodiversity Information Facility (GBIF), the GTDB r226 or the Microbe Atlas Project (see Methods).

**Extended Data Figure 8.**
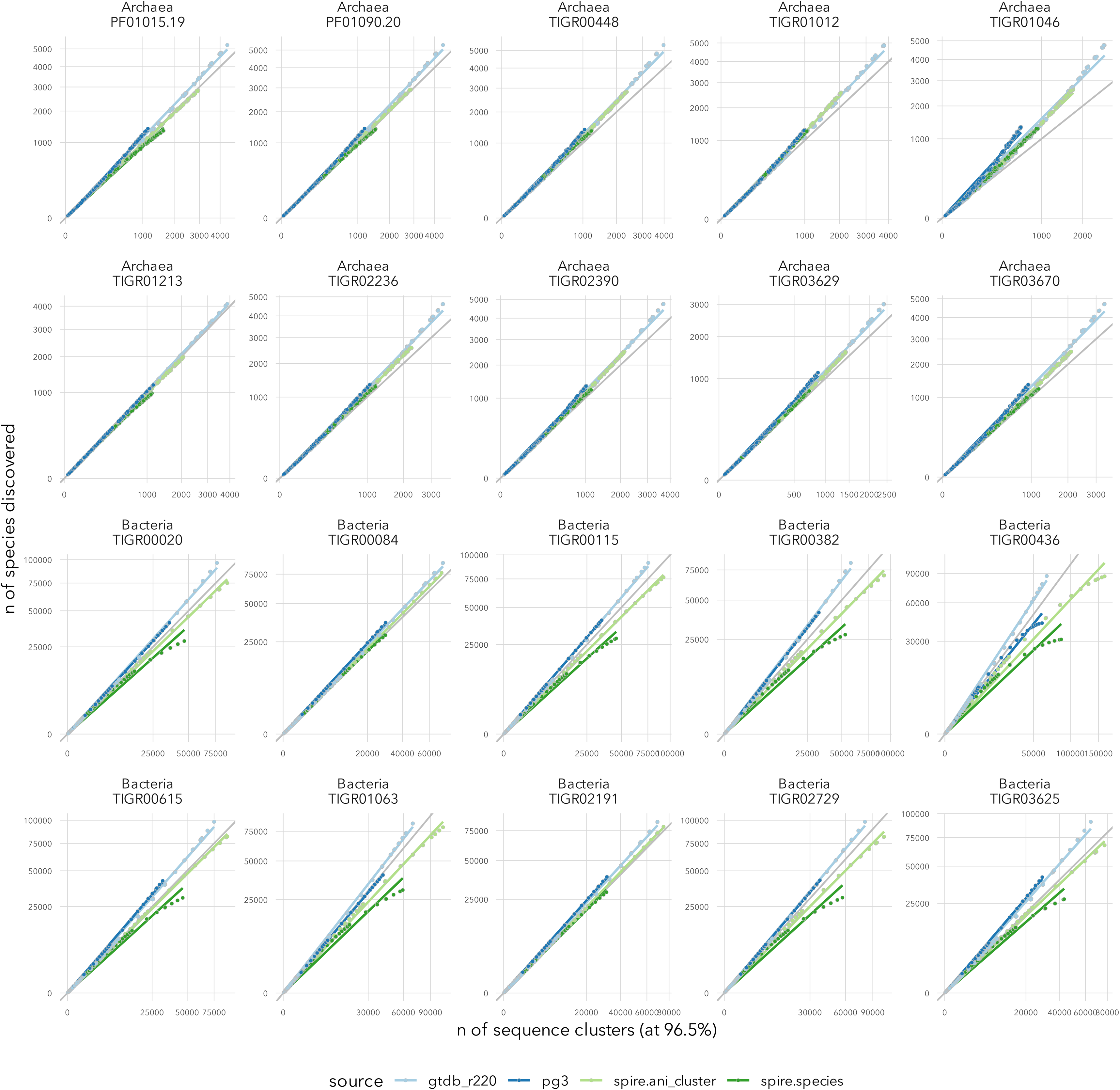
Marker gene-specific parameters to estimate species counts. Plots show the number of discovered species (y axis) against the number of sampled marker gene sequence clusters (x axis), for 10 randomly picked marker genes each for Archaea and Bacteria. The underlying data was obtained from iterative rarefaction along logarithmic steps (see Methods) for four different data series: ‘**GTDB r220’** refers to (clustered) marker genes originating from GTDB r220 reference genomes, and corresponding species definitions; ‘**pg3**’ are marker genes extracted from proGenomes3, and corresponding GTDB-based species classifications; ‘**spire.ani_cluster**’ uses SPIRE MAGs clustered at 95% ANI as ‘species’ clusters (i.e., analogous to GTDB definitions) and MAG-derived marker genes; ‘**spire.species**’ uses species-level GTDB classifications of SPIRE MAGs instead. The deviation of the ‘spire.species’ series at higher marker gene cluster counts is likely due to ‘underclassification’ (a significant fraction of SPIRE MAGs were not classifiable to species level against the GTDB reference). Therefore, in analyses shown in the main text, coefficients fitted from the ‘spire.ani_cluster’ series were used to estimate species counts from (unbinned) marker gene clusters.

**Extended Data Figure 9.**
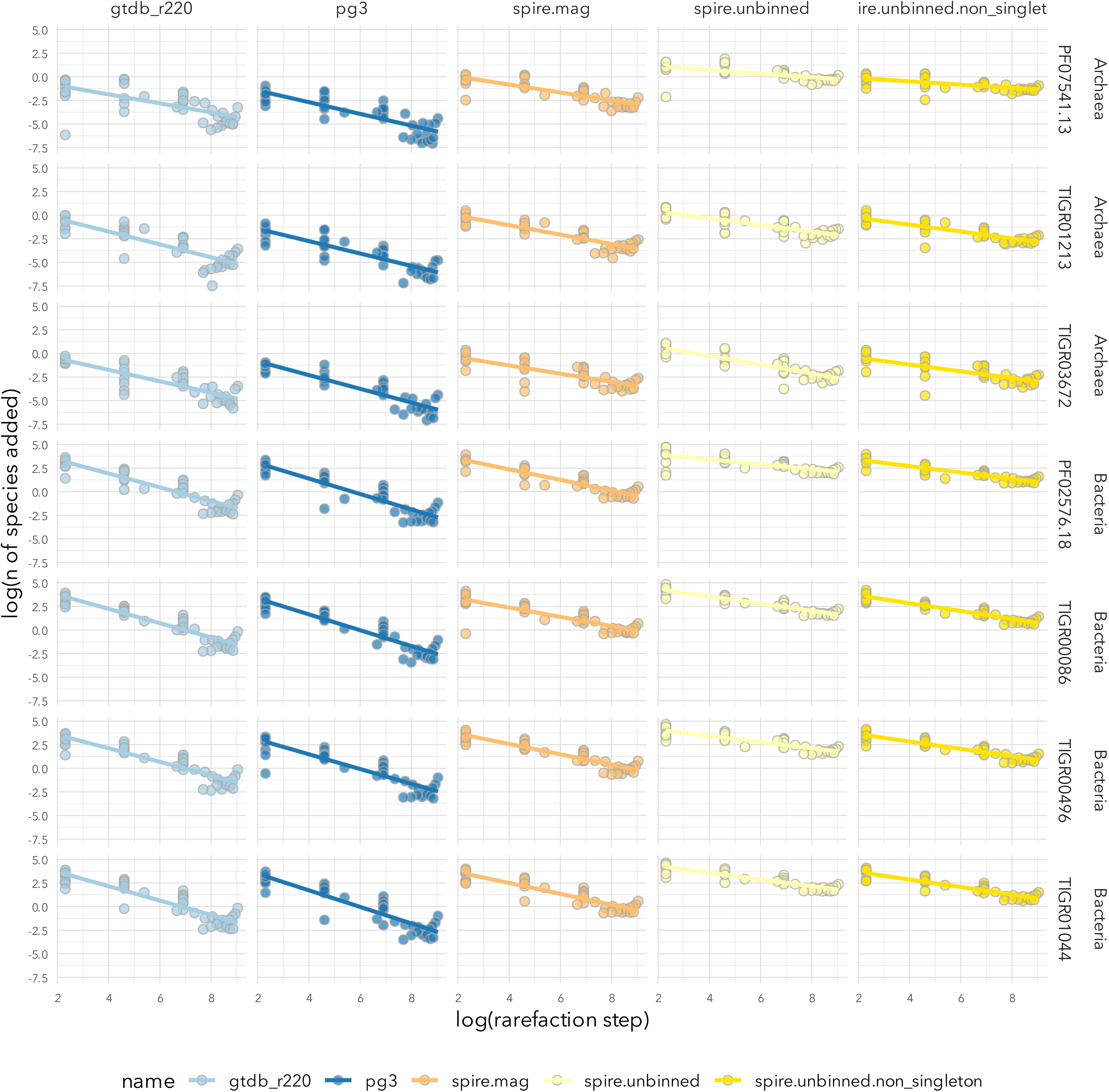
Fitting species discovery coefficients. Plots show the underlying data to fit ‘species discovery coefficients’ α (see Methods) for three randomly picked marker genes each for Archaea and Bacteria. Based on rarefactions, the log-log plots show the number of added species per sample (i.e., number of additionally ‘discovered’ species, y axis) per rarefaction step (i.e., number of samples added to the survey, x axis). Species discovery coefficients were estimated from the corresponding regressions in log-log space.

## Supplementary Figure Legends

**Figure S1. Habitat-stratified rarefaction curves for Archaea.** Species discovery curves as described in the main text and shown in Figure 1, but subset by individual habitats.

**Figure S2. Habitat-stratified rarefaction curves for Bacteria.** Species discovery curves as described in the main text and shown in Figure 1, but subset by individual habitats.

## Supplementary Table Legends

**Table S1. Definitions and data overview for microbial habitats considered in this study.** The table contains a full list of the various habitat categories, defined based on *microntology* annotations of samples (see Methods). It further provides a data overview including the number of samples per habitat and estimated species counts (across marker genes) in different data sources.

**Table S2. Sample metadata.** *microntology* annotations and assigned habitat category for each sample considered in this study.

**Table S3. Marker gene overview.** Number of extracted sequences per data source for each marker gene considered in this study.

