## Supplementary figures and images for "Hidden but discoverable diversity in the global microbiome"

### Supplementary Figure 1

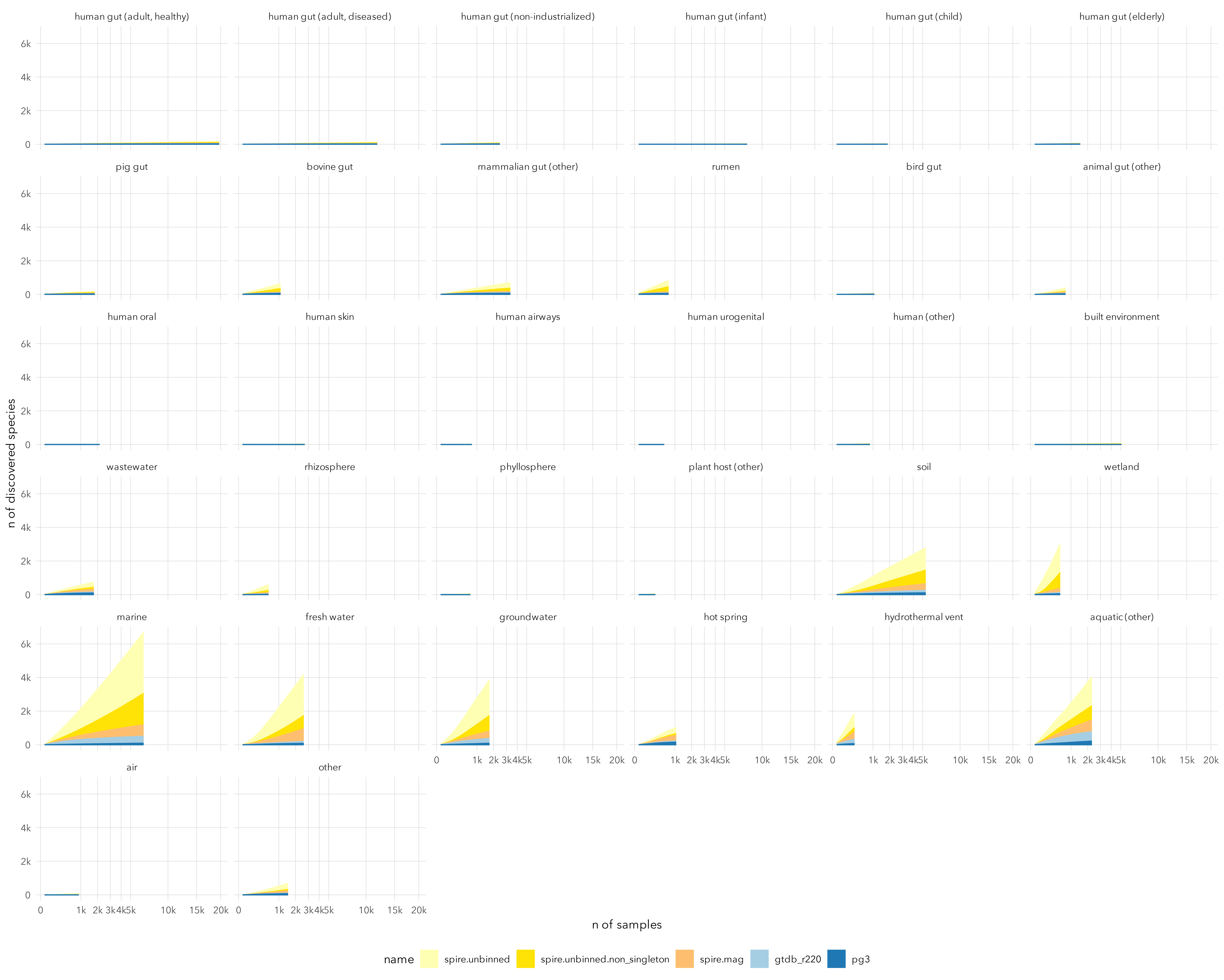

### Supplementary Figure 2

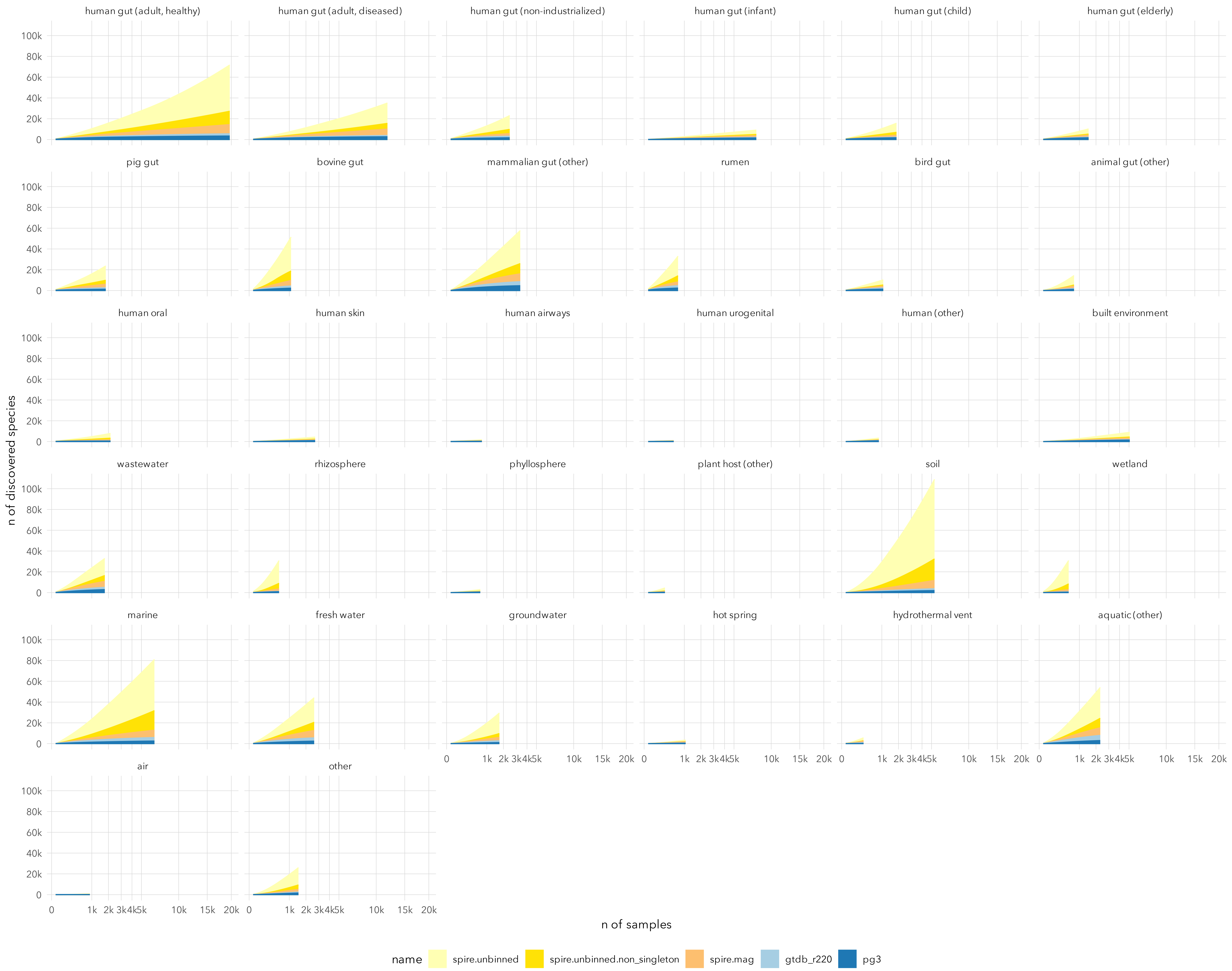
