## Supplementary Discussion for "Hidden but discoverable diversity in the global microbiome"

### Biological signal and noise among singleton marker gene clusters.

A pertinent question for all marker gene-based surveys of (microbial) diversity is *accuracy*. An ideal approach is both *sensitive* and *precise*: it recovers ‘real’ biological signal (i.e., sequence variants representing real biological lineages) with low rates of both false negatives (real sequences that are missed) and false positives (erroneous calls, due to noise introduced e.g. by sequence artifacts).

Amplicon-based surveys arguably provide only limited accuracy. Amplification bias can introduce false positives based on ‘chimeric’ artifacts, but the use of different primers (with differential taxonomic coverage) on different gene subregions introduces more complex skews [1]. Moreover, definitions of similarity cutoffs for ‘species-level’ resolution in 16S rRNA remain debated, and different lineages may indeed share identical amplified subsequences [2]. Finally, the focus on just one taxonomic marker prohibits links to functional or genomic context for uncultivated lineages.

An integrated survey across multiple marker genes based on metagenomic assemblies mitigates most of these issues, yet may introduce others. In particular, a major fraction of observed (species-level) marker gene clusters were *singletons*, i.e. containing only one sequence. Below, we discuss to what extent these singleton clusters may be due to noise (i.e., artifacts introduced during data generation and processing) and why we are confident that most indeed represent ‘real’ biological lineages.
**Possible assembly artifacts**. A recent benchmark estimated metagenomic misassembly errors to be in the range of ~7% of contigs [3]. Assembly errors are not randomly distributed and most are expected to be either (i) intra-species chimera (which would not greatly affect species-level clustering) or (ii) of such a type that HMMs would no longer detect a (partial) gene of sufficient length to pass our filters [4]. In a large-scale benchmark of ~300M assembled ORFs as part of the Global Microbial Gene Catalogue study [5], 75% of singleton genes were detectable in multiple metagenomic samples based on raw reads; 92% fell into known gene families; and only 0.4% were potential chimeras based on very inclusive criteria. We therefore estimate that misassembled genes with gene cluster-breaking errors are rare in our analysis.

**Singleton gene clusters are not disproportionately overrepresented among unbinned contigs.** Extended Figure 4 shows that the fraction of singleton clusters derived from MAGs and unbinned contigs is higher than for reference genomes (proGenomes3), but not disproportionately so. Indeed, a significant fraction of (singleton) clusters among reference genomes was also not detected among our metagenomic assemblies. In our analyses on clade size distributions (Willis / Yule curves, Figure 4), the estimated ‘rho’ values (Yule-Simon shape parameter) for Archaea and Bacteria are lower for SPIRE-derived data (including unbinned contigs) than for GTDB or GBIF (Global Biodiversity Information Facility) archaeal and bacterial reference taxonomies, as well as than for 16S-based OTUs from the Microbe Atlas Project. Among other things, lower ‘rho’ values correspond to size distributions that are less dominated by singletons and doubletons. In other words, SPIRE-derived clade size distributions are broadly in line with reference databases, but if anything, they are less heavy on singleton and small clades.

**Possible underestimation of diversity due to alternate genetic codes.** Our study does not explicitly account for alternative genetic codes; our ORF calls (and HMM searches) were conducted per metagenomic sample and gene callers perform poorly at detecting alternate codes (see e.g. [6] or [7]), so it is likely that a significant number of real ORFs were prematurely truncated which would lead to an overall under-estimation of diversity.

Taken together, we are confident that most singleton marker gene clusters in our data correspond to real biological entities, rather than technical or data artefacts, and that diversity estimates excluding singleton clusters represent a conservative lower bound.
